# Age, sex, and mitochondrial-haplotype influence gut microbiome composition and metabolites in a genetically diverse rat model

**DOI:** 10.1101/2024.10.28.620746

**Authors:** Hoang Van M. Nguyen, Eleana Cabello, David Dyer, Chloe Fender, Manuel Garcia-Jaramillo, Norman G. Hord, Steven Austad, Arlan Richardson, Archana Unnikrishnan

**Affiliations:** Department of Nutritional Sciences, College of Allied Health, University of Oklahoma Health Sciences, 1200 N Stonewall Ave, Oklahoma City, OK 73117, US. HVMN; Department of Microbiology and Immunology, College of Medicine, University of Oklahoma Health Sciences, Oklahoma City, OK 73117. US. EC,; DD; Environmental and Molecular Toxicology, College of Agricultural Sciences, Oregon State University, 2750 SW Campus Way, Corvallis, OR 97331, US. CF,; MGJ; Department of Nutritional Sciences, College of Education and Human Sciences, Oklahoma State University, 122 N Monroe St, Stillwater, OK 74075, US. NGH; Department of Biology, College of Arts and Sciences, University of Alabama at Birmingham, 902 14; Street South, Birmingham, AL 35205, US. SA; Department of Biochemistry and Physiology, College of Medicine, University of Oklahoma Health Sciences, 975 NE 10th Street, Oklahoma City, OK 73104, US. AR,, AU; Oklahoma Center for GeroScience and Healthy Brain Aging, University of Oklahoma Health Sciences, 975 NE 10th Street, Oklahoma City, OK 73104, US. AR; Oklahoma Veteran Affairs Medical Center, Oklahoma City, Oklahoma, 921 NE 13th St, Oklahoma City, OK 73104, US. AR; Harold Hamm Diabetes Center, OU Health, Oklahoma City, Oklahoma, 1000 N Lincoln Boulevard, Oklahoma City, OK 73104, US. AU

**Keywords:** Gut microbiome, gut microbial community, aging, mitochondria, rat, metabolomics

## Abstract

We evaluated the impact of sex and mitochondrial-haplotype on the age-related changes in the fecal gut microbiome of the genetically heterogeneous rodent model, the OKC-HET^B/W^ rat. Alpha-diversity, measuring richness and evenness of gut microbiome composition, did not change with age or mitochondrial-haplotype. However, beta-diversity, a measure of microbial differences among samples, was significantly modulated by age in male and female rats in both mitochondrial-haplotypes. The age-related changes in the microbiome differed markedly between male and female rats. Five microbial species changed significantly with age in male rats compared to nine microbial species in female rats. Only three of these microbes changed with age in both male and female rats. The mitochondrial-haplotype of the rats also affected how aging altered the microbiome. Interestingly, most of the microbial species that changed significantly with age were mitochondrial-haplotype and sex specific, i.e., changing in one sex and not the other. We also discovered that sex and mitochondrial-haplotype significantly affected the age-related variations in content of fecal short-chain fatty acids and plasma metabolites that influence or are regulated by the microbiome, e.g., tryptophan derived metabolites and bile acids. This study demonstrates that the host’s sex plays a significant role in how the gut microbiome evolves with age, even within a genetically diverse background. Importantly, this is the first study to show that the mitochondrial-haplotype of a host impacts the age-related changes in the microbiome and supports previous studies suggesting a bidirectional interaction between the gut microbiome and host mitochondria.

**Highlights:** Most age-related changes in microbial species occurred in one sex but not the other

Mitochondrial-haplotype altered the microbiome and was generally sex dependent

Microbiome associated metabolites differed by age, sex, and mitochondria-haplotype

## 1. Introduction

Since Lane et al. developed an efficient technique to sequence the 16S rRNA gene [1, 2], characterization of commensal microbes has illuminated the pleiotropic effects of the microbiome on health and disease. For example, studies have shown that the fecal gut microbiome (herein described as microbiome) influences the host’s metabolic health [3], immunity [4], cardiovascular health [5], and cognitive function [6, 7]. These effects in the host have drawn the attention of aging researchers because dysbiosis, or perturbations in microbiome composition, is currently considered a hallmark of aging [8]. A review by Badal et al. [9] shows that the human microbiome changes with age. However, there is little consensus on how age and the microbiome interact in humans because age-related changes in the host’s diet [10] and environment [11] can have a major impact on microbe abundance. These confounding variables make it difficult to draw conclusions on how aging specifically affects the microbiome in human studies. Use of laboratory rodents, where the diet and environment can be controlled throughout the life of an animal, allow investigators to directly test the impact of the aging host on its microbiome.

Studies in rodents support data in humans and show that the microbiome changes with age in mice [12–15] and rats [16–18]. In addition, studies with mice have shown that aging interventions that increase lifespan and healthspan, such as caloric restriction or rapamycin, affect the age-related changes in the microbiome [19, 20]. However, several gaps in our knowledge must still be addressed to gain a better understanding of how the microbiome changes as the host ages. One problem is that many of the past studies in rodents only assess age-related changes of the microbiome in one sex in mice [12–15, 21] or rats [16–18]. Sex differences with age have also been largely ignored in human studies, with Badal et al. [9] reporting that 8 out of 27 papers did not disclose sex of their participants and 15 out of 27 studies combined male and female participants for statistical analysis. Data in humans and mice show that serum sex hormone levels can alter microbiome composition [22–24]. In addition, sex hormone levels decrease with age with the changes occurring differently in male and female humans and rodents [25–29]. Another limitation in the current studies with mice is that all have used inbred animals, which have negligible genetic variation. Because the genotype of the host can impact the microbiome [30–33], the lack of genetic variation in most rodent studies not only limits our knowledge of how aging affects the microbiome in other rodent genotypes but also limits the translation of the data from rodents to humans.

In this study, we have used a novel OKC-HET^B/W^ rat model to study the impact of age and sex on the microbiome in a genetically heterogenous animal model. In addition, this rat model allows us to determine for the first time if mitochondrial-haplotype (mt-haplotype), which has been shown to impact gut microbiome composition in young, inbred mice [34–36], has an impact on the age-related changes in the microbiome. We found that the abundance of various microbial species changed significantly with age in the genetically heterogenous OKC-HET^B/W^ rats. Importantly, most of the age-related changes in the microbiome were both sex and mt-haplotype dependent. In addition, we observed that the changes in the microbiome were associated with changes in short-chain fatty acids in the feces and in microbiome derived metabolites in the host’s plasma.

## 2. Materials and Methods

### 2.1 Animals

The OKC-HET^B/W^ rats with two different mitochondrial haplotypes were generated by breeding four inbred strains of rats [(Brown Norway (BN), Fischer 344 (F344), Wistar Kyoto (WKY), and Lewis (LEW) rats] obtained from Charles River as previously described [37]. Briefly, BN/F344 F1 rats were generated by crossing female BN rats to male F344 rats and WKY/LEW F1 rats were generated by crossing female WKY to male LEW rats (WKY/LEW). OKC-HET^B^ rats were generated by crossing female BN/F344 to male WKY/LEW rats and the OKC-HET^W^ were generated by crossing female WKY/LEW to male BN/F344 rats. Due to this selective breeding and taking advantage of the maternal inheritance of mitochondrial DNA, all rats have similar nuclear heterogeneity while containing two different mitochondrial haplotypes. The OKC-HET^B^ rats contain mitochondria from the BN rats, and the OKC-HET^W^ rats contain mitochondria from the WKY rats, which differ by 94 nucleotides [37]. These rats were bred and maintained in the Oklahoma City VA Medical Center animal facilities in specific pathogen free conditions. They were fed *ad libitum* on chow diet (Picolab Rodent Diet 5053, LabDiet, St. Louis, MO). Male and female OKC-HET^B^ and OKC-HET^W^ rats were studied at 9- (adult) and 26- (old) months of age. Feces and plasma were collected from adult male OKC-HET^B^ (n=6), adult male OKC-HET^W^ (n=6), adult female OKC-HET^B^ (n=9), adult female OKC-HET^W^ (n=9), old male OKC-HET^B^ (n=10), old male OKC-HET^W^ (n=6), old female OKC-HET^B^ (n=7), and old female OKC-HET^W^ (n=8) rats. Rats were fasted for 16 hours prior to termination and whole blood was collected in EDTA coated tubes by cardiac puncture, and plasma was separated from whole blood (1000xg for 10 minutes), flash frozen, and stored at -80°C. The colon was separated from the anus and feces was collected directly from the colon, immediately flash frozen in liquid nitrogen, and stored at -80°C. All procedures were approved by the Institutional Animal Care and Use Committee at the Oklahoma City Veterans Affairs Health Care System.

### 2.2 Microbiome Analysis

#### 2.2.1 DNA Extraction

DNA was isolated from colon fecal samples using the ZymoBIOMICS DNA miniprep Kit (D4300, ZYMO RESEARCH, Orange, CA) as specified by the manufacturer. A NanoDrop Lite Spectrophotometer (ThermoFisher Scientific) was used to determine quantity and quality of isolated DNA.

#### 2.2.2 16S rRNA Sequencing

Library construction and 16S rRNA sequencing were performed on the isolated fecal DNA by the OUHS Institutional Research Core Facility (IRCF). Data were generated using Illumina MiSeq libraries prepared using MiSeq Reagent Kit V3-V4. Data analysis was provided by the OK-INBRE Data Science Core. Sequences were processed and analyzed using QIIME2 v2022.11 [38]. Standard data clean-up was performed using Cutadapt [39] Sequences were grouped into amplicon sequence variants (ASVs) using the DADA2 QIIME2 plugin [40]. MAFFT was used to align the ASVs and FastTree was used to create a rooted phylogenetic tree [41, 42]. Rarefaction curves showed that all samples reached asymptote indicating the sequencing depth used was sufficient. A QIIME2 naïve Bayesian classifier trained on sequences from the V3-V4 region of Greengenes v13_8 99% OTUs was used to assign a taxonomic profile to each ASV [43]. Microbial abundance tables were generated to the species level.

### 2.3 Analysis of Plasma Metabolites

#### 2.3.1 Metabolite Extraction

Extraction of metabolites from rat plasma was adapted from a previous study [44]. Metabolites from 70 µL of plasma were extracted in 400 µL cold methanol/acetonitrile (1:1, v/v) and homogenized in a Precellys 24 Touch Homogenizer (Bertin technologies, Montigny-le-Bretonneux, France) twice for 20 seconds. The samples were then incubated for 2 hours at -20°C to precipitate protein. Samples were centrifuged at 4°C for 15 minutes at 13,000xg, and 400 µL of supernatant was collected. The supernatant was centrifuged again at 4°C for 15 minutes at 13,000xg. Supernatant was collected and evaporated to dryness in a vacuum concentrator for 2 hours. The dry extracts were reconstituted in 150µL of acetonitrile/DI water (1:1, v/v) and then vortexed for 30 seconds to dissolve the dried metabolite pellet. The samples were centrifuged for the last time at 4°C for 10 minutes at 13,000xg. A quality control (QC) pooled sample was prepared by combining 5 µL of each sample supernatant. The QC is a “mean” profile representing all metabolites encountered in the analysis. The sample supernatants and QC were stored at -80°C until they were analyzed using liquid chromatography with tandem mass spectrometry (LC-MS/MS).

#### 2.3.2 LC-MS/MS-based Metabolomics Analysis

Samples were spiked just before analysis with 3 µL of isotope labeled metabolite Mix 1 QReSS Kit (Cambridge Isotope Labs) to account for potential instrument performance variation throughout the analysis. **Supplementary** Figure 1. shows that our range of variation between the expected m/z and measured m/z was less than 0.05 ppm. Liquid chromatography was performed using a Sciex ExionLC AD ultra-high-performance liquid chromatograph (UPHLC) system. Volumes of 2 µL were injected into a 2.1 x 150 mm, 2 µL Intersil Ph-3 HPLC column (GL Sciences, Torrance, CA). The autosampler and column over temperature were held at 15 °C and 40 °C, respectively. A binary elution system of (A) LC-MS grade water + 0.1 % formic acid and (B) methanol + 0.1% formic acid was utilized to achieve separation using a flow rate of 0.3 mL/min in a 23 min gradient. Pooled QC and blanks were injected between every 8 samples, and samples were fully randomized prior to injection. Mass spectrometry analysis was conducted with a Sciex ZenoTOF 7600 (Framingham, MA) in both positive and negative electrospray ionization modes, utilizing the information-dependent acquisition (IDA) mode.

#### 2.3.3 LC-MS/MS data processing

Raw data was imported into MS Dial v5 (RIKEN Center, Yokohama City, Kanagawa, Japan) [45] for feature detection, peak alignment, and peak integration. Metabolites were confirmed using MS, MS/MS fragmentation using publicly available libraries for LC-MS/MS from the MassBank of North America (MoNA), and an in-house curated IROA library (Ann Arbor, Michigan). Data was acquired in both electrospray ionization (ESI) positive and negative modes. First, features were removed from the dataset if their peak height was not at least 5-fold higher than that found in blank samples. Features without MS/MS peaks, or known fragmentation signatures, were removed. Metabolites associated with microbial deconjugation of bile acids (primary and secondary bile acids) and tryptophan metabolism (serotonin, kynurenine, indoles) were selected from the metabolome for further data analysis [46–48].

### 2.4 Targeted Metabolomics for Fecal Short-chain Fatty Acids (SCFAs)

Colonic feces (200 mg) of five randomly selected rats per group were sent to Metabolon (Metabolon, INC., NC) where SCFAs were quantified with a targeted metabolomic LC-MS/MS (Agilent 1290 UHPLC/Sciex QTrap 5500) panel of eight total metabolites. All eight metabolites were detected in our samples.

### 2.5 Statistical Analysis

Statistical analysis was completed using MicrobiomeAnalyst 2.0 [49], MetaboAnalyst 6.0 [50], and GraphPad v10 (Dotmatics, Boston, Massachusetts). Statistical analysis of male and female rats was completed separately. Comparisons of age (adult vs. old) were performed in rats of the same sex and mt-haplotype. Comparisons of mt-haplotype were performed on rats of the same age and sex but different mt-haplotypes (e.g. adult male OKC-HET^B^ vs. adult male OKC- HET^W^). ANOVA with Tukey’s post-hoc analysis was used to determine differences in body weight, subcutaneous fat weight, and gonadal fat weight. T-test was used to compare fat weights in old female OKC-HET^B/W^ rats.

MicrobiomeAnalyst 2.0 was used to determine differences in Chao1 and Shannon alpha-diversity using Mann-Whitney/Krushkal-Wallis FDR<0.05 cutoff for significance. Differences in beta-diversity were determined using Jensen-Shannon Divergence and PERMANOVA FDR<0.05 to determine significance and were represented by principal coordinate analysis (PCoA) plots. Differences in abundance of individual microbes at different levels of taxa were determined using Mann-Whitney/Kruskal-Wallis (MK) FDR<0.05 and Linear Modeling (LM) from MaAsLin2 [51] integrated in MicrobiomeAnalyst 2.0 after normalizing data using relative log expression. For ease of understanding, all significant differences are represented as relative abundance (%).

Statistical analysis of metabolite content was performed in GraphPad v10. Heat maps were created using MetaboAnalyst 6.0. Comparisons were completed in male and female rats separately. Fischer’s LSD with FDR < 0.05 was used to determine significance. Additionally, differences in metabolites were considered to be marginally significant when Fischer’s LSD <0.5, and FDR >0.05. This was done to bring attention to metabolites that could be physiologically important though they did not meet the arbitrary significance value of p=0.05. All variables were checked for normality and non-parametric tests were utilized if the test for normality was significant (i.e. the data were not normal).

## 3. Results

### 3.1 Changes in fecal gut microbiome composition with age differ by sex and mt-haplotype in OKC-HET^B/W^ rats

To examine how sex and mt-haplotype affect the age-related changes in gut microbiome composition, we used a rat model our group developed using a four-way cross strategy with four commercially available inbred rat strains (BN, F344, LEW, and WKY) as described in the Methods. Two F2 lines were created that were heterogenous with respect to the nuclear genome that had one of two mt-haplotypes: mitochondria from either BN (OKC-HET^B^) or WKY (OKC- HET^W^) rats. Rat body weights and composition are shown in **Supplementary** Figure 2. Male rats had increased body weight with age. While subcutaneous and gonadal fat tended to increase with age, the increase was not statistically significant. Female rats showed an increase in body weight and the weight of subcutaneous and gonadal with age. Interestingly, the subcutaneous and gonadal fat weight were significantly increased in the old female OKC-HET^W^ rats compared to the OKC-HET^B^ rats. After identifying commensal gut microbes based on the 16S rRNA sequence in these rats, our dataset contained 205 operational taxonomic units (OTUs). OTUs that appeared in only one sample were removed to reduce artifacts, leaving 149 features for the fecal samples from male and female rats.

We first measured alpha-diversity of the microbiome using both the Chao1 and Shannon Index to test for differences in OTU richness and evenness. There were no significant differences in either the Chao1 or Shannon Index when comparing male or female rats by age or mt-haplotype (**Figure 1)**. However, the Chao1 index was marginally significant in adult female OKC-HET^B^ compared to the adult female OKC-HET^W^ and old female OKC-HET^B^ suggesting decreased richness in adult female OKC-HET^B^ rats.

**Figure 1.**
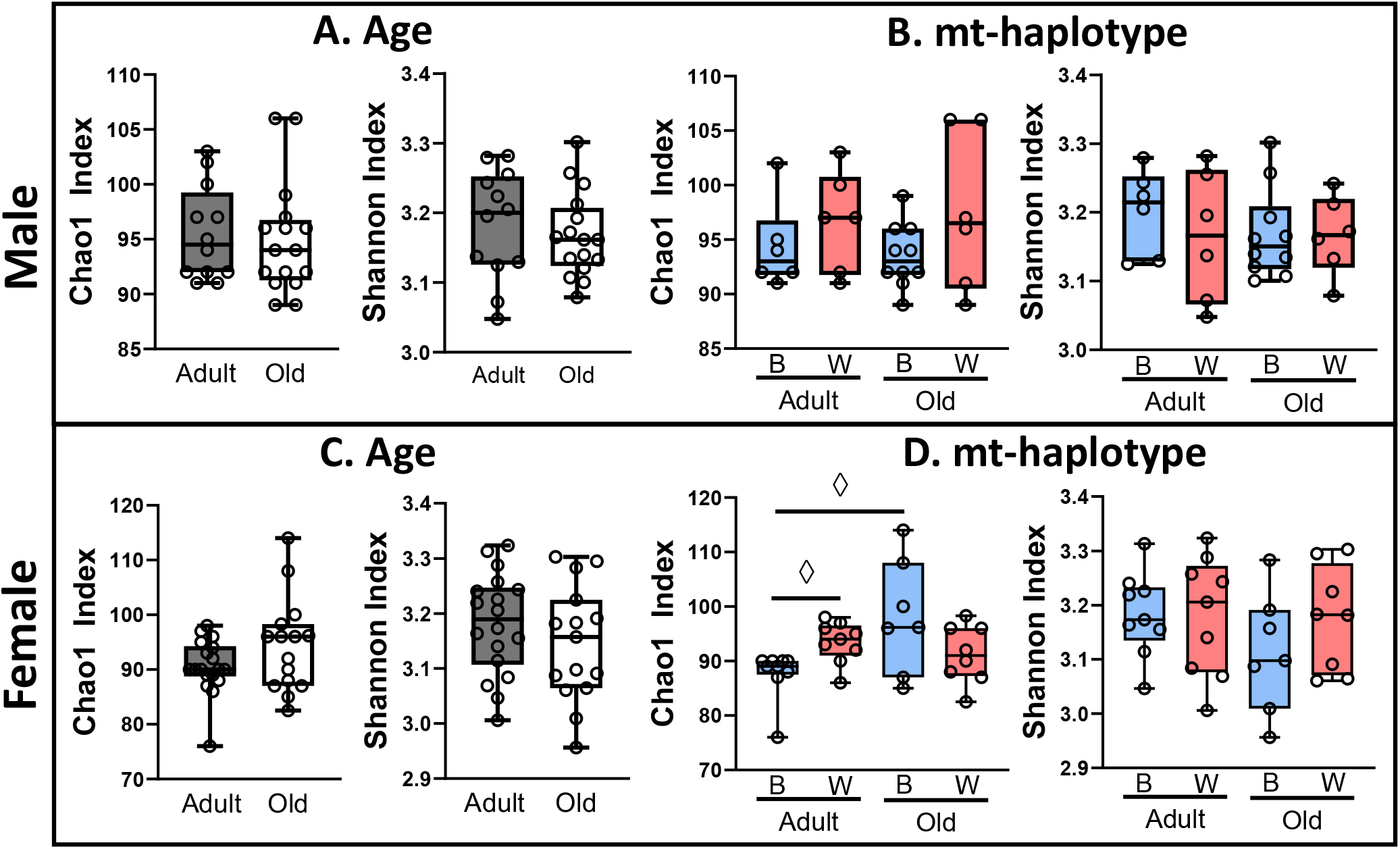
Alpha-diversity is not significantly different with age and mt-haplotype in OKC-HET^B/W^ rats. Alpha-diversity of the microbiome from adult (9-months) and old (26- months) OKC-HET^B/W^ rats was measured by the Chao1 and Shannon index. The effect of age on alpha-diversity is shown for adult (gray boxes) and old (white boxes) male (A) and female (C) rats with the two mt-haplotypes combined. The effect of the OKC-HET^B^ (blue boxes) and OKC-HET^W^ (red boxes) haplotype on alpha-diversity is shown for male (B) and female (D) rats. The data were collected from 6 to 10 rats per group, and the box plots display the 1^st^ and 3^rd^ quartiles with a horizontal line at the median. The whiskers display minimum and maximum values. ^◊^Values marginally significant by Fischer’s LSD p < 0.05.

Next, we evaluated how the overall microbiome composition changed in male and female rats using beta-diversity; a distance-based analysis of similarities or dissimilarities between samples. Beta-diversity was visualized using principal coordinate analysis (PCoA). The PCoA plots in **Figure 2** show age comparisons (adult vs. old) in rats of the same sex and mt-haplotype. In male rats, the microbiome composition was significantly different by age in both OKC-HET^B^ (**Figure 2A**) and OKC-HET^W^ rats (**Figure 2B)** . A similar pattern was observed in female rats where beta-diversity was significantly different by age in OKC-HET^B^ (**Figure 2C**) and OKC- HET^W^ rats (**Figure 2D**). Differences in the microbiome composition by mt-haplotype (OKC- HET^B^ vs. OKC-HET^W^) was also evaluated using beta-diversity in rats of the same age and sex. There were no significant differences in beta-diversity when comparing mt-haplotype in either male or female rats (**Figure 2E-H**).

**Figure 2.**
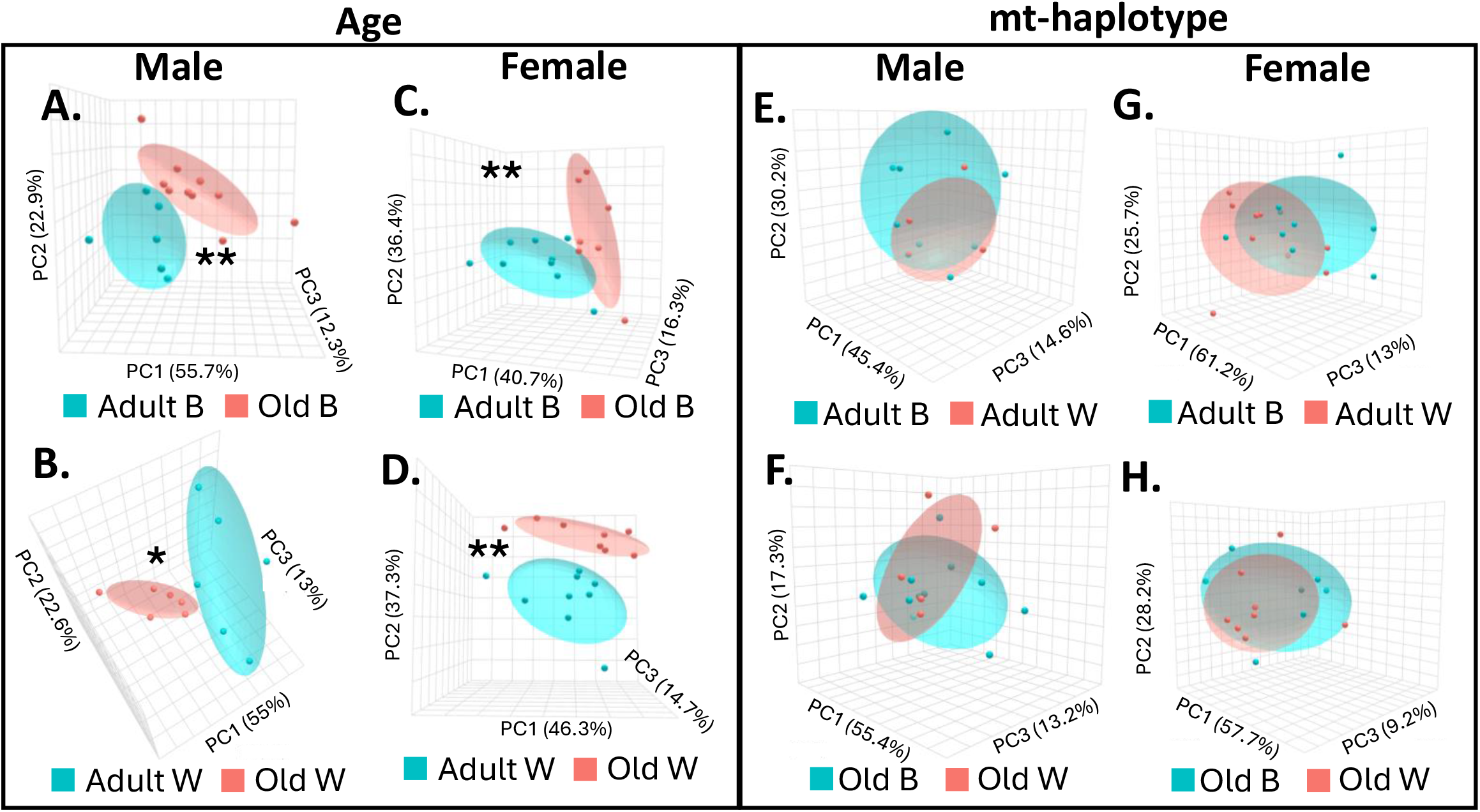
Beta-diversity is different with age in OKC-HET^B/W^ rats. Beta-diversity of the microbiome was measured using Jenssen-Shannon divergence comparing adult (9-months, blue ovals) and old (26-months, red ovals) rats for the following: male OKC-HET^B^ (A), male OKC- HET^W^ (B), female OKC-HET^B^ (C), and female OKC-HET^W^ (D) rats. Beta-diversity was also compared between OKC-HET^B^ (blue ovals) and OKC-HET^W^ (red ovals) haplotypes for the following: old male (F), adult female (G), and old female (H) rats. Oval outlines represent the 95% confidence interval for each group. The data were collected from 6 to 10 rats per group, and those values statistically significant by PERMANOVA at p<0.05* and p<0.01** are shown.

Given the significant differences in beta-diversity by age, we next measured the abundance of microbial species that changed with age for male and female OKC-HET^B^ and OKC-HET^W^ rats. The identifiable species (45 species for male, 47 species for female) in our dataset accounted for 0.83-0.96% of the relative abundance for male, and 0.81-0.97% relative abundance in females. Most of the species that changed significantly with age were sex and mt-haplotype specific except for *R. callidus*, which decreased in both males and females (**Figure 3**). In males, the abundance of five microbial species were identified that changed significantly with age (**Figure 3A**). *R. callidus* abundance decreased with age in both mt-haplotypes. *L. reuteri* increased, *R. albus* decreased, and *L. garvieae* decreased in male OKC-HET^B^ rats but not OKC- HET^W^ rats. *C. saccharogumia* abundance was significantly increased in male OKC-HET^W^ but not in male OKC-HET^B^ rats.

**Figure 3.**
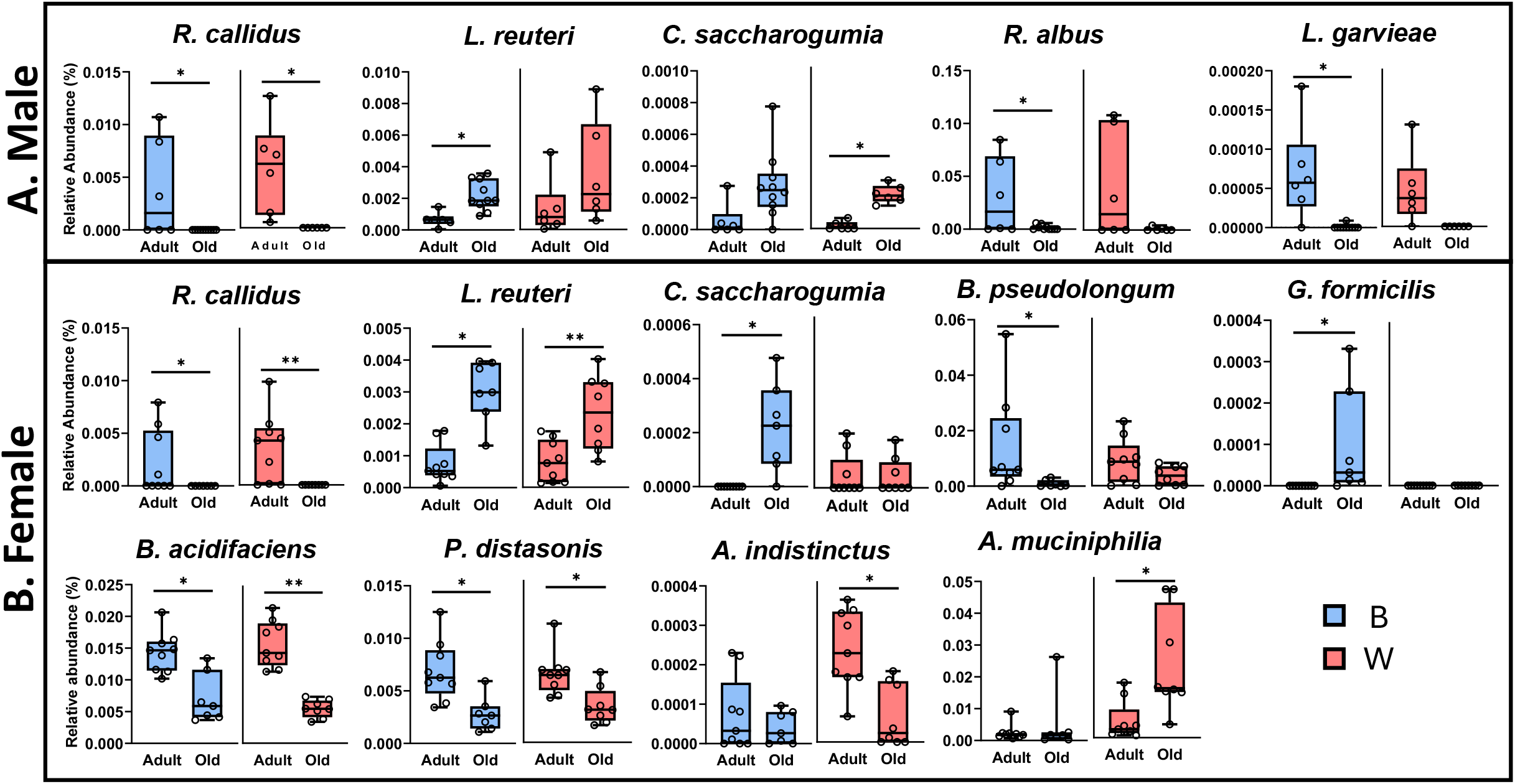
Microbial species that are significantly changed with age are dependent on sex and mt-haplotype. Microbial species in the gut microbiome from male (A) or female (B) rats were compared for adult (9-months) and old (26-months), OKC-HET^B^ (blue boxes) and OKC-HET^W^ (red boxes) rats. The box plots display the 1^st^ and 3^rd^ quartiles with the horizontal line denoting the median, and the whiskers display minimum and maximum values. The data were collected from 6 to 10 rats per group and statistically compared using Mann-Whitney/Kruskal-Wallis and/or Linear Modeling with significance at p<0.05*, p<0.01** shown.

In female OKC-HET^B^ and OKC-HET^W^ rats, we observed significant changes in the abundance of nine microbial species (**Figure 3B**), three of which also changed in male rats (*R. callidus, L. reuteri,* and *C. saccharogumia)*. The abundance of *R. callidus, B. acidifaciens,* and *P. distasonis* decreased while *L. reuteri* increased with age in both female OKC-HET^B^ and female OKC-HET^W^ rats. In addition, the abundance of *C. saccharogumia* and *G. formicilis* increased while *B. pseudolongum* decreased with age in female OKC-HET^B^ but not female OKC-HET^W^ rats. In female OKC-HET^W^ rats, *A. indistinctus* abundance decreased and *A. muciniphilia* increased with age. In total, the abundance of four species of microbes changed significantly with age in male OKC-HET^B^ while only 2 species changed significantly with age in male OKC- HET^W^ rats. In female rats, the abundance of a similar number of microbe species changed in the two mt-haplotypes (e.g., 7 species for OKC-HET^B^ rats and 6 species for OKC-HET^W^ rats).

We next compared the abundance of microbes at every level of taxonomy by mt-haplotype in rats of the same sex and age (**Figure 4**). We did not identify any significant differences at the species level. However, we observed that the abundance of genus *Lachnospira* was significantly increased in adult male OKC-HET^W^ rats compared to OKC-HET^B^ (**Figure 4A**) rats. In female rats, the abundance of genus *Bilophila* was significantly increased in the adult OKC-HET^W^ rats compared to adult OKC-HET^B^ rats (**Figure 4B**). In old female rats, a significant increase in the abundance of the order *Verrucomicrobiales* was observed in OKC- HET^W^ rats compared to OKC-HET^B^ rats (**Figure 4B**). In our dataset, the genera *Lachnospira* and *Bilophila* are individual OTUs and do not have species level data. *A. mucinipilia* is the only microbial species associated with the order *Verrucomicrobiales* in our dataset.

**Figure 4.**
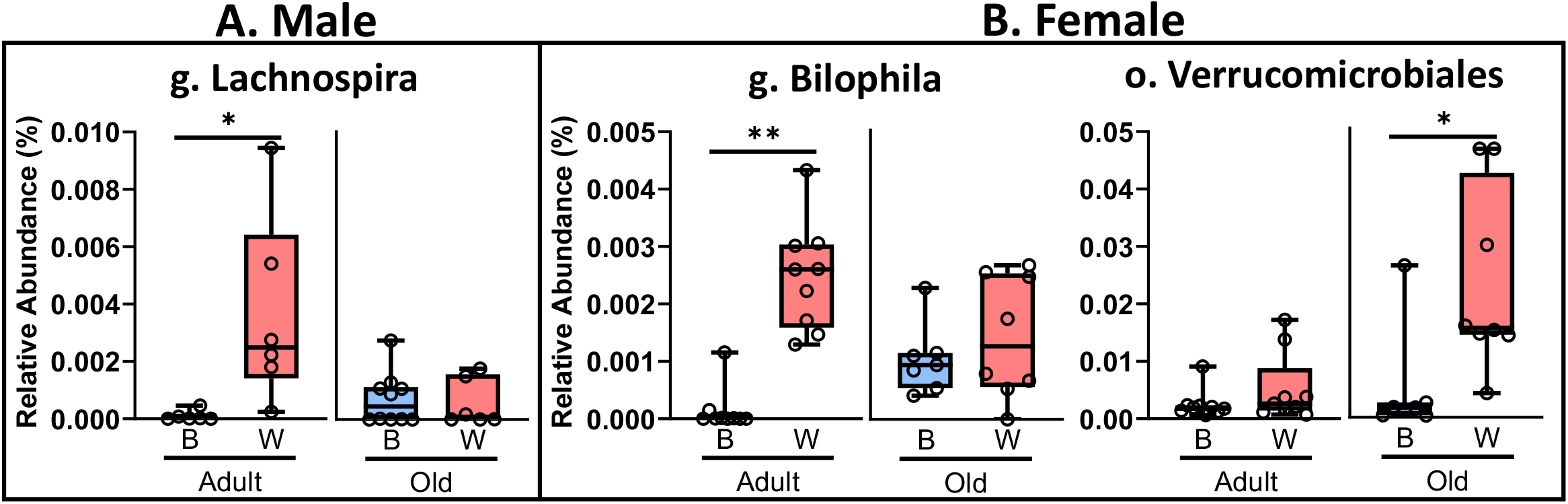
The mt-haplotype affects several microbe groups in OKCHET^B/W^ rats. Microbial OTUs are compared for OKC-HET^B^ (blue boxes) and OKC-HET^W^ (red boxes) male (A) and female (B) rats. The box plots display the 1^st^ and 3^rd^ quartiles with a horizontal line the median. The whiskers display minimum and maximum values. The data were collected from 6 to 10 rats per group, and the mt-haplotypes were statistically compared using Mann-Whitney/Kruskal- Wallis and/or Linear Modeling with significance at p<0.05*, p<0.01** shown.

### 3.2 Changes in fecal short-chain fatty acids (SCFAs) with age differ by sex in the OKC- HET^B/W^ rats

SCFAs, such as butyric acid, acetic acid, and propionic acid, are generated from microbial fermentation of fiber. SCFAs have been shown to impact the host by regulating metabolic pathways in the host [52]. Therefore, we measured the abundance of SCFAs in the feces collected from the colon of our rats to determine if there were changes in SCFAs produced by the microbiome with age and mt-haplotype. **Supplementary** Figure 3 shows a heat map of the average normalized content of the eight SCFAs we detected in the fecal material. These heat maps of SCFAs profile suggested an increase in fecal SCFA in old OKC-HET^W^ rats regardless of sex. However, we only observed a significant age-related increase in total SCFAs in female OKC-HET^W^ rats and a marginal increase in female OKC-HET^B^ rats (**Supplementary** Figure 3D). **Figure 5** shows the three SCFAs that changed significantly (FDR q-value <0.05) or were marginally significant (Fisher’s LSD p-value<0.05 and FDR q-value>0.05) in female rats. In female rats, the increase in acetic acid was significant for both OKC-HET^B^ and OKC-HET^W^ rats when comparing age. Propionic acid levels increased with age in female OKC-HET^W^ rats, while the levels of hexanoic acid tended to increase with age in female OKC-HET^W^ rats. There were no significant changes in SCFAs when comparing mt-haplotypes in male or female rats.

**Figure 5.**
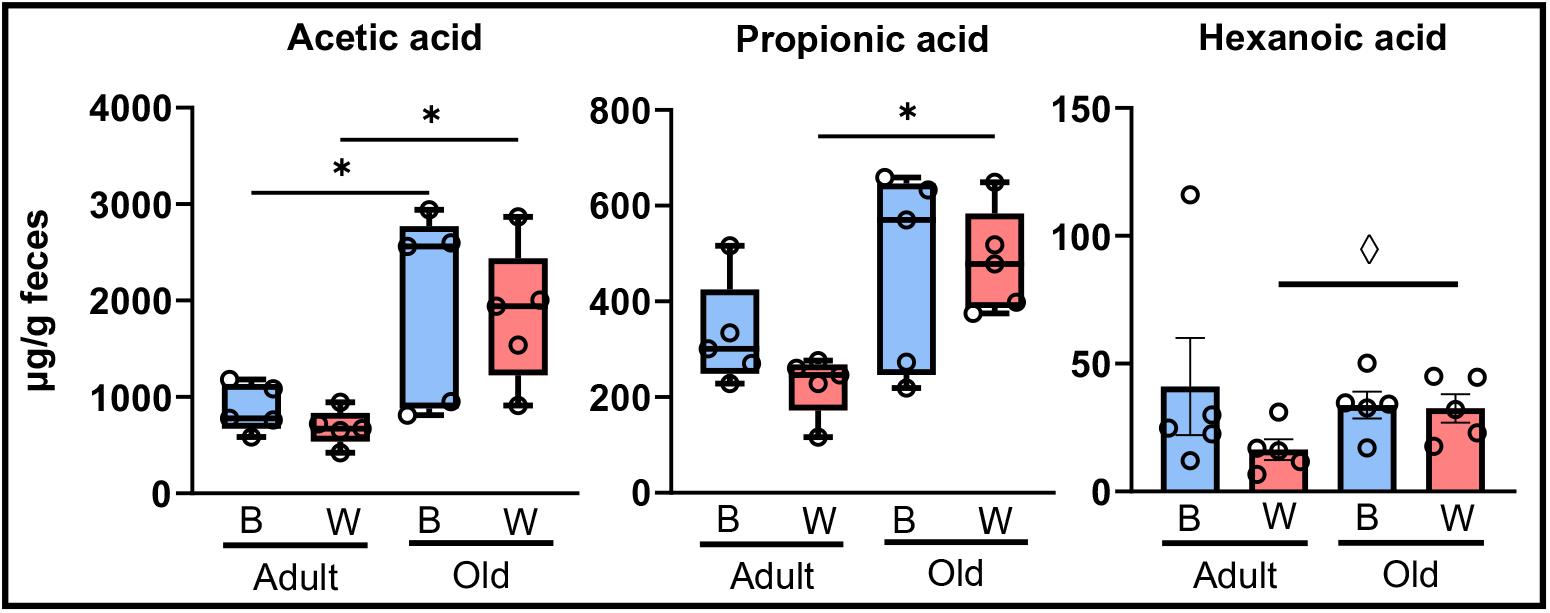
Fecal short chain fatty acids changed with age in female rats. Fecal SCFAs were measured in adult (9-months) and old (26-months), OKC-HET^B^ (blue boxes) and OKC-HET^W^ (red boxes) rats. Only female rats, shown here, were found to exhibit a significant difference with age in fecal SCFAs. The box plots display the 1^st^ and 3^rd^ quartiles with the horizontal line the median. The whiskers display minimum and maximum values. The data were collected from 5 rats per group, and values significantly different by FDR q<0.05* or values marginally significant by Fischer’s LSD p < 0.05^◊^ are shown.

### 3.3 Changes in plasma tryptophan and bile acid metabolites with age differ by sex and mt-haplotype in the OKC-HET^B/W^ rats

Gut microbes are known to communicate with the host via microbe derived plasma metabolites. Therefore, we measured the various tryptophan and bile acid metabolites in the plasma using an untargeted metabolomics dataset, which was obtained from the same rats that were used to study the microbiome. Dietary tryptophan is an essential amino acid that is metabolized by microbes into bioactive compounds. **Supplementary** Figure 4 shows a heat map of the average levels of the nine tryptophan metabolites identified in the plasma isolated from OKC-HET^B^ and OKC-HET^W^ rats. These metabolites arise from the three major arms of tryptophan metabolism: 1) indoles, 2) serotonin production, and the 3) kynurenine pathway. The most striking difference between the mt-haplotypes were observed in the indole profile of female rats which suggested that female OKC-HET^W^ rats had reduced plasma levels of the indole metabolites compared to the OKC-HET^B^ rats regardless of age (**Supplementary** Figure 4D).

Figure 6 shows the levels of the plasma tryptophan metabolites that changed significantly (FDR q-value <0.05) or were marginally significant (Fisher’s LSD p-value<0.05) with age or mt-haplotype in male and female rats. In male rats, there were no tryptophan metabolites that changed significantly when comparing age or mt-haplotype; however, kynurenine was marginally increased in adult male OKC-HET^W^ rats compared to OKC-HET^B^ rats (Figure 6A). In contrast, female rats showed a significant change with age and mt-haplotype in several tryptophan metabolites (Figure 6B). Adult female OKC-HET^B^ rats had higher plasma content of kynurenine and hydroxykynurenine compared to old female OKC-HET^B^ rats. Additionally, indoxyl-3-sulfate levels were marginally increased with age in female OKC-HET^B^ rats. Only kynurenine levels decreased significantly with age in female OKC-HET^W^ rats. When comparing female rats by mt-haplotype, kynurenine and hydroxykynurenine plasma levels were significantly increased in the adult female OKC-HET^B^ compared to their OKC-HET^W^ counterparts, while the increase in 5-hydroxytryptophan levels were marginally significant in adult female OKC-HET^W^ compared to adult female OKC- HET^B^ rats. Interestingly, mt-haplotype differences only occurred in adult animals.

**Figure 6.**
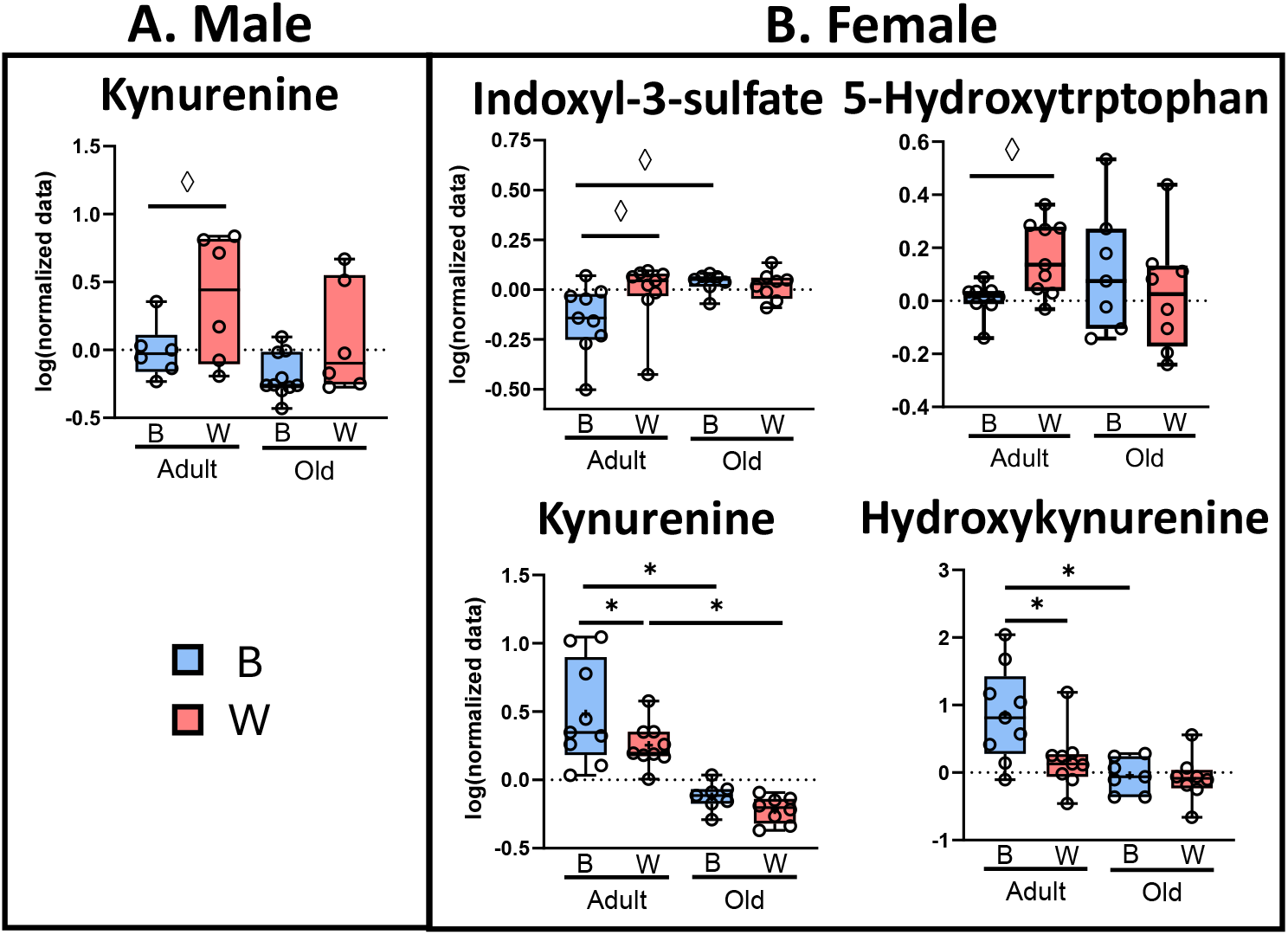
Microbial metabolism of tryptophan to kynurenine may be affected by age and mt-haplotype in female rats. Plasma tryptophan metabolites from male (A) and female (B) OKC-HET^B^ (blue boxes) and OKC-HET^W^ (red boxes) adult (9-months) or old (26-months) rats are shown. The box plots display the 1^st^ and 3^rd^ quartiles and a horizontal line at the median, and the whiskers display minimum and maximum values. The data were collected from 5 randomly selected rats per group, and the values significantly different by FDR q<0.05* or values marginally significant by Fischer’s LSD p < 0.05^◊^ are shown.

Bile acids are another mode of host-microbiome communication. Primary bile acids produced from cholesterol are conjugated with either taurine or glycine in the liver and released into the gastrointestinal tract in rats. Approximately, 95% of the bile acids are recycled through enterohepatic circulation. However, ∼5% of the primary bile acids reach the colon and are metabolized by the gut microbiome to produce secondary bile acids and can be absorbed to facilitate host-microbiome communication [53].

**Supplementary** Figure 5 shows a heat map of the average plasma content of the fourteen bile acids and taurine that were detected in the plasma of the rats. In male rats, the overall bile acid profile suggests a robust increase in primary bile acids in plasma with age in both male OKC-HET^B^ and OKC-HET^W^ animals that was not apparent in female rats. Figure 7 shows the plasma content of the seven bile acids that changed with age and/or mt-haplotype. In male rats (Figure 7A), there was a significant increase in the levels of tauroursodeoxycholic acid (TUDCA) while the levels of taurochenodeoxycholic acid (TCDCA), glycocholic acid (GCA), and glycoursodeoxycholic acid (GUDCA) were marginally significant in male OKC-HET^B^ rats with age but not in OKC-HET^W^ rats. Cholic acid (CA) was the only bile acid that significantly increased with age in male OKC-HET^W^ rats. When comparing mt-haplotype in male rats, cholic acid levels were significantly increased in old male OKC-HET^W^ rats compared to their OKC- HET^B^ counterparts. In female rats (Figure 7B), the change in bile acids was limited to two metabolites with only plasma levels of lithocholytaurine (LCT) showing a significant decrease with age in both OKC-HET^B^ rats and OKC-HET^W^ rats (Figure 7B). The levels of taurodeoxycholic acid (TDCA) levels were marginally increased in old female OKC-HET^W^ rats compared to adult OKC-HET^W^ rats and old female OKC-HET^B^ rats.

**Figure 7.**
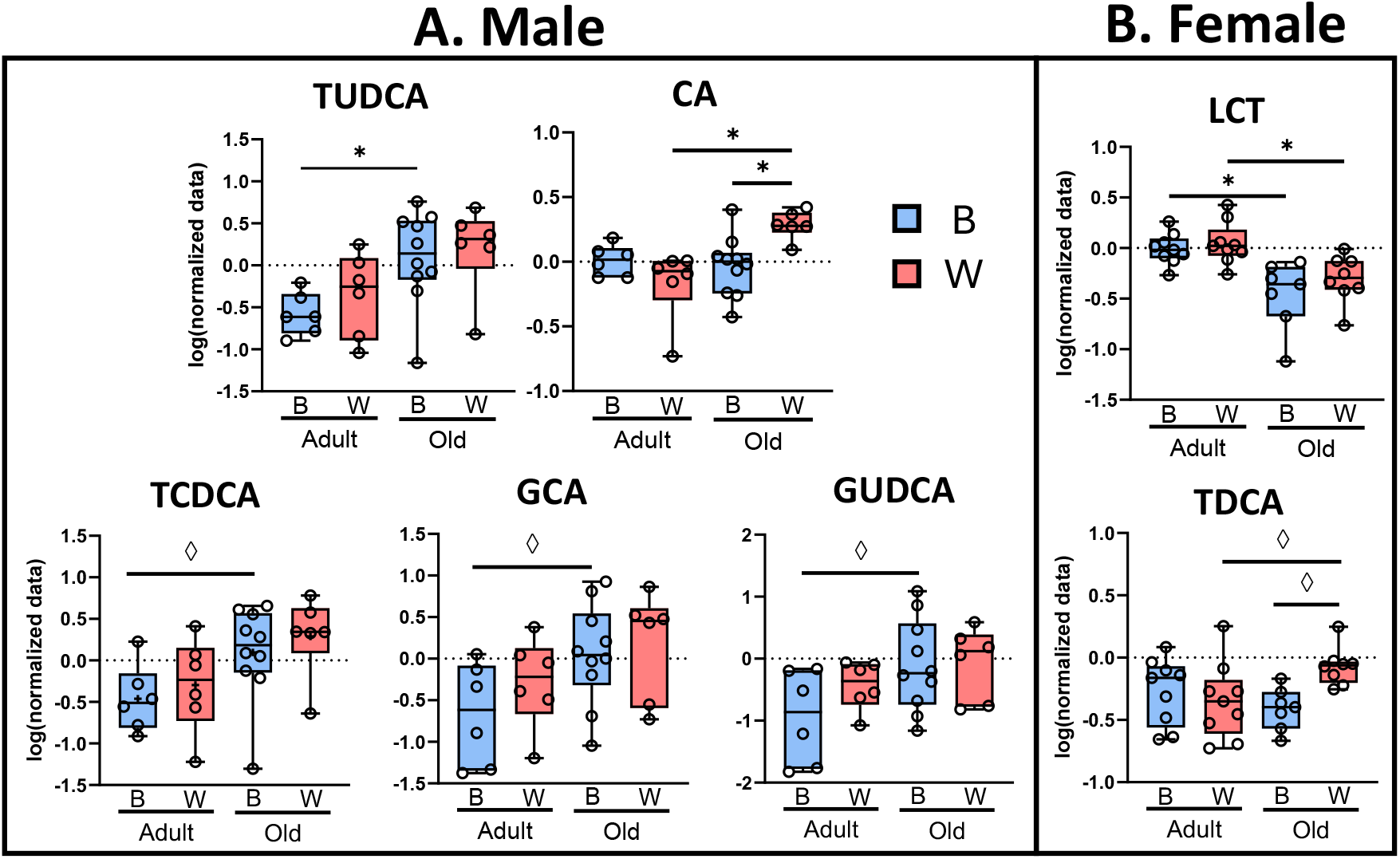
Changes in primary, but not secondary bile acids, are dependent on sex and mt-haplotype. Plasma bile acids from male (A) and female (B) OKC-HET^B^ (blue boxes) and OKC-HET^W^ (red boxes) adult (9-months) or old (26-months) rats are shown. The box plots display the 1^st^ and 3^rd^ quartiles with the horizontal line for the median, and the whiskers display minimum and maximum values. The data were collected from 6 to 10 rats per group, and significance was defined as FDR q<0.05* or values marginally significant by Fischer’s LSD p < 0.05^◊^. TUDCA, tauroursodeoxycholic acid; CA, cholic acid; TCDCA, taurochenodeoxycholic acid; GCA, glycocholic acid; GUDCA, glycoursodeoxycholic acid; LCT, lithocholytaurine; TDCA, taurodeoxycholic acid

## 4. Discussion

Over the past two decades, investigators have studied how aging impacts the gut microbiome. Laboratory rodents have emerged as an excellent model to study microbiome changes with age because variables that can affect the microbiome composition in humans, such as diet and environment [9], can be controlled over the lifespan of the animal. One of the limitations in previous rodent studies is that they have used inbred mouse strains [12–15, 21]. Because the genome of the host has been shown to affect the gut [31–33], it is unknown how well microbiome changes with age in one inbred mouse strain will translate to another inbred strain or to a genetically diverse population, such as that encountered by investigators in studying humans. This problem is exacerbated because all aging studies in mice have used only one inbred strain, C57BL/6 mice. While several studies have used outbred rat strains, e.g., Wistar or Sprague Dawley rats [16–18], the genetic variation is limited in these outbred rats [54] and can vary considerably from one commercial source to another [55]. To overcome these limitations, we used a genetically heterogenous rodent model, the OKC-HET^B/W^ rat, which was generated by a four-way cross using four commercial inbred strains of rats that were deliberately selected to maximize genetic heterozygosity [37]. This breeding strategy not only allows us (or other investigators) to generate similar genetically heterozygous rats at any time; but also allows us to generate two strains of rats that differ in their mitochondrial genomes, e.g., mitochondria from either BN (OKC-HET^B^) or WKY (OKC-HET^W^) rats. Several reports (Hirose et al., Kunstner et al., and Yardeni et al.) indicate that mt-haplotype can alter the microbiome of young C57BL/6 mice [34–36]. Thus, the OKC-HET^B/W^ rat gives us the first opportunity to determine if the mt-haplotype has an impact on the age-related changes in the microbiome in a genetically heterozygous animal model. Another advantage of using the OKC-HET^B/W^ rat model is that rats are more similar to humans in many respects than laboratory mice used in research, including the prevalence of age-related pathologies [56], which could impact the microbiome in old animals. Therefore, the data generated from the OKC-HET^B/W^ rats are likely more translatable to humans.

Unfortunately, most of the aging studies on the gut microbiome in humans and rodents have either used only one sex (almost always males), failed to disclose the sex, or combined males and females into one group. Therefore, we studied the impact of age and mt-haplotype on the microbiome in both male and female OKC-HET^B/W^ rats. We found that beta-diversity, which is a measure of distance-based dissimilarities between-samples and depicted with PCoA plots (Figure 2), showed a clear separation by age in both male and female OKC-HET^B/W^ rats. The age-related changes in beta-diversity agree with reports in which beta-diversity has been measured in C57BL/6 mice [13–15], Sprague-Dawley rats [16], and humans [9].

To determine how the gut microbiome changed with age, we compared the abundance of microbial species in adult (9 months) and old (26 months) male and female OCK-HET^B/W^ rats. A total of eleven microbial species changed significantly with age: five species in male rats and nine species in female rats (Figure 3). Only three microbes, *R.callidus, L.reuteri,* and *C. saccharogumia* changed significantly with age in the same direction in both male and female rats, e.g., *R. callidus* decreased with age and *L. reuteri, C. saccharogumia* increased. The two species that changed with age in only male rats were *R. albus* and *L. garviea* which both decreased. On the other hand, six species changed with age only in females. *A. mucinipilia* and *G. formicilis* increased with age and *B. psuedolongum, B. acidifaciens, A. indistinctus,* and *P. distasonis* decreased. Thus, we found that the sex of the host played an important role in shaping the microbiome changes that occur with age in the OKC-HET^B/W^ rat. Currently, there are only two studies that have compared the effect of sex on the age-related changes in the microbiome: one in rats [57] and one in humans [58]. Lee et al. studied the effect of feeding a high-fat diet to young rats (6 weeks) and old (2 years) inbred F344 rats [57]. While they found that feeding a high-fat diet significantly altered the microbiome of young and old male and female rats, they did not find any significant age-related change in relative abundance of microbes in either male or female chow fed F344 rats. The limited number of rats studied (especially the old rats) and/or the collection of fecal material after defecation, which can impact the microbiome composition [59], could explain the differences in our findings. Takagi et al. studied the gut microbiome in healthy

Japanese subjects ranging from 20 to 89 years of age [58]. There were no significant changes in the microbiome with age observed at any level of taxonomy in either males or females separately. However, significant differences were observed in the microbiome between male and female participants at each age group studied. In fact, fourteen genera were increased specifically in males compared to females, and eleven genera were significantly increased in female compared to male participants. While it is clear that sex plays a major role in gut microbiome composition, our data are the first to demonstrate that sex of the host had a major impact on the age-related changes in the gut microbiome and emphasizes the importance of studying both males and females when investigating the impact of aging on the microbiome.

The OKC-HET^B/W^ rat model allowed us to study the effect of the mt-haplotype on the age-related changes in the gut microbiome for the first time. Previous studies have reported that the mitochondrial genome of the host could affect the microbiome of young, inbred mice where male and female mice were grouped together for analysis [34–36]. However, these studies are limited because the mitochondrial genomes compared differed by only 2 to 5 nucleotides, which is a result of the traditional inbred strains of laboratory mice originating from a single female *Mus musculus domesticus* mouse [60]. In addition, the different mitochondrial genomes were on the same inbred background (C57BL/6). In contrast, the mitochondrial genomes of the OKC- HET^B^ and OKC-HET^W^ rats differ by 94 nucleotides, which is much more similar to the differences in the mitochondrial genomes seen in humans [61]. Importantly, because the B- and W-haplotypes are on a genetically heterogenous background, any differences we observe in the OKC-HET^B^ and OKC-HET^W^ rats are robust, i.e., changes occur on multiple nuclear genotypes and are therefore more likely to occur in humans. We observed that most of the microbial species that changed significantly with age occurred in one but not both mt-haplotypes. For example, nine out of eleven microbes that changed with age were mt-haplotype specific and occurred in one sex and not the other. *R. callidus* was the only microbial species we found that changed with age in both male and female rats and in both mt-haplotypes. In male rats, four other microbial species (*L.reuteri, C. saccharogumia, R. albus, L.garvieae*) changed with age in one mt-haplotype but not the other. In female rats, *L.reuteri, B. acidifaciens,* and *P. distasonis* changed with age in both OKC-HET^B^ and OKC-HET^W^ rats. Five other microbial species *(C. sacchargoumia, B. pseudolongum, G. formicilis, A. indistinctus, A. mucinipilia*) changed in one mt-hyplotype and not the other. Although the previous studies did not show any association between the mt-haplotype and microbial species, they reported significant associations at the genus [34, 35] and family [36]. We also found mt-haplotype differences at the level of the genus and order. For example, the genus *Lachnospira* was increased in adult male OKC-HET^W^ compared to adult male OKC-HET^B^ rats. The family *Lachnospiraceae* (containing genus *Lachnospira*) was reported to differ with mt-haplotypes in C75BL/6 mice [34, 36] and was negatively correlated with reactive oxygen species production [36]. We found the genus *Bilophila* was significantly increased in adult female OKC-HET^W^ rats compared to adult female OKC-HET^B^ rats. The family *Desulfovibrionaceae*, containing the genus *Bilophila*, was reported to be significantly increased in C57BL/6J mice with the FVB/NJ mouse mt-haplotype compared to their conplastic pairs containing C57BL/6 nuclear DNA and NZB/BlnJ mt-haplotype [34]. Finally, we observed that the order *Verrucomicrobiales* was significantly increased in old female OKC-HET^W^ compared to old female OKC-HET^B^. These observed differences when comparing mt-haplotypes show that mt-haplotype can influence microbiome composition in a genetically heterogenous animal model.

To determine how the changes in the microbiome might impact the aging host, we measured the SCFAs. SCFAs are produced from fermentation of indigestible fiber from commensal bacteria in the gut and can regulate metabolic pathways in the host [52]. We observed a trend for an age-related increase in total fecal SCFAs in female rats with female OKC-HET^W^ rats having a significant increase in total SCFAs with age. This change with age can be attributed to increased acetic acid in old female OKC-HET^B^ and OKC-HET^W^ rats and increased propionic acid in old female OKC-HET^W^ rats. Previous studies in humans have reported that fecal SCFAs decrease with age in both male and female human participants [62] and when male and female participants are combined [63]. Additionally, Lee et al. [57] found that cecal SCFAs decreased with age in male F344 rats. However, fecal SCFAs were observed to increase in obese compared to lean individuals [64] and was positively correlated with risk factors associated with metabolic syndrome in women such as increased adiposity [65]. Therefore, we compared the level of total fecal SCFAs to the fat mass of the individual female rats. As shown in **Supplementary** Figure 6, we observed a positive correlation between total fecal SCFAs and subcutaneous fat mass that tended towards significance (p=0.07, r=0.41) in female rats. When separating female OKC- HET^W^ rats, there was a significant positive correlation (p=0.04, r=0.65). The changes in SCFAs in feces could arise from an increase in microbes that produce SCFAs [66, 67] or from decreased absorption of SCFAs into plasma [68]. Because most of the bacteria known to produce SCFAs, e.g., Lactobacillaceae and Ruminococcaceae families [52], decreased with age in our rats (Figure 3), the increased SCFAs we observed in feces of the old female rats most likely arises from a decrease in the absorption of the SCFAs into the plasma, which could arise from the age-related decline in intestinal absorption that has been reported in rats [69], mice [70], and humans [71].

Tryptophan metabolites are one of the best examples of the metabolic cross talk between the gut microbiome and host because these metabolites can influence the host’s health and disease processes [46, 72]. Tryptophan is metabolized by microbes into indoles or modulates tryptophan metabolism through pathways producing either serotonin or kynurenine. The production of serotonin and kynurenine are primarily produced by the host’s endochromaffin cells [73, 74] and liver [75, 76], respectively. However, indole and its derivatives are produced specifically by gut microbiota [46]. We observed major differences in plasma metabolites of tryptophan by sex. The total level of plasma indoles tended to be lower in adult and old female OKC-HET^W^ rats compared to OKC-HET^B^ rats (**Supplementary** Figure 4D). Four tryptophan metabolites changed with age in female rats and only one in males (Figure 6). The plasma levels of kynurenine showed the greatest change with age in females. Although the mechanism through which the microbiome impacts liver kynurenine production is unclear, studies using germ-free mice have shown that plasma levels of kynurenine are significantly reduced in the absence of the microbiome [75], which is increased after recolonization [77]. Plasma kynurenine decreased significantly with age in both mt-haplotypes in female rats. Although kynurenine levels did not change significantly with age in male rats, they were marginally decreased in male OKC-HET^B^ compared to male OKC-HET^W^ rats. A study in male inbred mice reported age-related increases in serum kynurenine levels between 3 and 28 months of age [15]. However, Comai et al. [78] reported an age-related decrease in enzymatic activity of liver indoleamine 2,3-dioxygenase (IDO) in Sprague-Dawley rats. IDO is the rate-limiting step in tryptophan metabolism to kynurenine and is modulated by the gut microbiome [46], which could agree with our observation that kynurenine levels decrease with age in rats. In a review. Bakker et al. reported that 50% of the human studies, which measured plasma kynurenine with age, found increased plasma kynurenine levels [79]. Interestingly, for all the tryptophan metabolites that changed with age, we observed mt-haplotype differences in adult rats. For example, kynurenine and hydroxykynurenine levels were reduced in female OKC-HET^W^ rats while in male rats, kynurenine was increased in male OKC-HET^W^ rats. Thus, mt-haplotype appears to play a role in tryptophan metabolism, which could occur via modulation of gut microbiome composition.

The microbiome also plays an important role in the bile acid pool size and composition [80]. Primary bile acids (PBAs) produced in the liver are secreted into the gastrointestinal tract of rat. Because primary conjugated bile acids are detergents and acidic, they can affect the gut microbe diversity and composition [47, 81]. While most of the PBAs are reabsorbed in the terminal ileum, the gut microbiota can deconjugate and metabolize them into secondary bile acids, which are absorbed in the terminal ileum and colon [80]. These secondary bile acids produced by the microbiome play an important role in bile acid homeostasis of the host, which can contribute to conditions like non-alcoholic fatty liver disease, inflammatory bowel disease, and cholesterol disorders [47, 82, 83]. We observed significant changes in the plasma levels of only PBAs with age and mt-haplotype. In contrast to tryptophan metabolites, the age-related changes in plasma levels of bile acids were greatest in male rats, with five metabolites changing with age or mt-haplotype compared with two metabolites in female rats. In general, there was an age-related increase in plasma bile acids in male but not female rats (**Supplementary** Figure 5). However, total primary bile acids were marginally significance with age only in male OKC- HET^B^ rats. These changes can be attributed to increases in the plasma levels of TUDCA, TCDA, GCA, and GUDCA in OKC-HET^B^ male rats. Our data support previous work in male rats suggesting increased plasma primary bile acids with age. For example, an increase in plasma bile acids was reported in male Wistar-Imamichi rats between 3 and 11 months of age [84], and taurine-conjugated bile acids were increased in bile collected from the bile duct between 6 and 15, months of age in male Sprague-Dawley rats [85]. We also observed that mt-haplotype had an impact on the plasma levels of the primary bile acids. Except for LCT, which decreased with age in both the B- and W-haplotypes in female rats, all the changes in bile acid metabolites were mt-haplotype specific. Particularly striking was the age-related increase in TUDCA, TCDCA, GCA, and GUDCA, which was observed only in male OKC-HET^B^ rats. We did not observe any differences in secondary bile acids when comparing age or mt-haplotype in male and female OKC-HET^B/W^ rats. Thus, age and mt-haplotype appear to play a greater role in the host’s ability to produce PBA with age than the plasma secondary bile acids produced by the microbiome. PBAs are known to influence gut microbiome composition because of their antimicrobial properties and by increasing the acidic environment of the colon [47, 81]. Therefore, we were interested in determining if any microbes changed specifically with age only in male OKC-HET^B^ rats that might be associated with the increase in plasma levels of TUDCA, TCDCA, GCA, and GUDCA. A significant increase in the abundance of *L. reuteri* was observed in old male OKC- HET^B^ rats that was associated with the age-related increase with PBAs in plasma of the male OKC-HET^B^ rats. Because *L*. reuteri is a lactobacillus species that is resistant to an acidic environment [81], the age-related increase in *L. reuteri* in the male OKC-HET^B^ rats might have arisen from increased PBAs in the colon that came from the increased levels of plasma PBAs.

In conclusion, we used a genetically heterogenous rat model to study how the gut microbiome changes with age that is more translatable to humans than previous studies with inbred mice or outbred rats, which have limited genetic diversity. We found the age-related changes in the microbiome differed greatly between male and female rats, demonstrating the importance of studying both males and females when evaluating the impact of age on the microbiome. Importantly, we found that mt-haplotype of the rats played an important role in how aging altered the microbiome. Although previous studies have shown the mitochondrial genome can affect the microbiome of young, inbred mice [34–36], it was not clear from these studies if the effect of mt-haplotype would translate to other genetic backgrounds or differ with sex. Our data show for the first time that mt-haplotype differences are robust enough to impact the microbiome on a genetic heterogenous background. In addition, we found that the effect of the mt-haplotype was sex dependent, i.e., the impact of the mt-haplotype on the age-related changes in the microbiome almost always occurred in one sex and not the other.

Recent data suggest a bidirectional interaction between the gut microbiome and mitochondria [75, 86, 87]. For example, an early study by Han et al. (2017) with *C. elegans* showed that several *E. coli* mutants promoted longevity through the secretion of colonic acid, which regulated mitochondrial dynamics and the unfolded protein response in the host’s cells. In addition, two species of *Lactobacillus* [44] and postbiotics from *Lacticaseibacillus casei* [88] were reported to alter mitochondrial function in the liver of rats. On the other hand, a deficiency of the mitochondrial protein (methylation-controlled J protein) in mice was shown to have profound effect on the microbiome [86, 89]. Thus, the question emerges as to how the microbiome and host mitochondria communicate. In a review, Zhang et al. (2022) proposed that SCFAs produced by the microbiome could be modulators of mitochondria function in the intestinal epithelial [87]. Interestingly, we showed that fecal SCFAs were increased with age in female rats. Yardeni et al. (2019) showed that differences in the mitochondrial redox status and ROS production that occurred in mice with different mitochondrial genomes were associated with modifications in the gut microbiome [36]. They proposed that changes in redox status might impact metabolites produced by the host cells that were then secreted into the gut. They also showed that expressing catalase in the mitochondria of the host had the single greatest impact of the gut microbiome, suggesting hydrogen peroxide produced by the host might act directly on the microbes in the gut. They also proposed that the release of mtDNA from the mitochondria in stressed intestinal cells could activate the cGAS-Sting pathway, resulting in an inflammatory response that could in turn affect the gut microbiome. Thus, to understand the impact of mt-haplotype on the gut microbiome, future studies should focus on how mitochondrial function differs in the cells in the intestine of OKC-HET^B^ and OKC-HET^W^ rats as they age. Our preliminary data show that feeding a high-fat diet affected mitochondrial function in skeletal muscle differently in the B- and W-haplotypes [37].

## CRediT Authorship

**HVMN –** conceptualization, data curation, formal analysis, investigation, methodology, visualization, writing – original draft, writing – review and editing; **EC** – data curation, methodology, writing – review and editing; **DD** – data curation, methodology, writing – review and editing; **CF** – data curation, methodology, writing – review and editing; **MGJ** – data curation, methodology, writing – review and editing; **NGH –** conceptualization, funding acquisition, writing – review and editing; **SA** – funding, methodology, writing – review and editing **AR** – supervision, funding acquisition, methodology, writing – original draft, writing – review and editing; **AU** – supervision, funding acquisition, methodology, writing – original draft, writing – review and editing.

## References

[1] D.J. Lane, B. Pace, G.J. Olsen, D.A. Stahl, M.L. Sogin, N.R. Pace, Rapid determination of 16S ribosomal RNA sequences for phylogenetic analyses, Proc Natl Acad Sci U S A 82(20) (1985) 6955–9.

[2] I.B. Zhulin, Classic Spotlight: 16S rRNA Redefines Microbiology, J Bacteriol 198(20) (2016) 2764–5.

[3] Y. Fan, O. Pedersen, Gut microbiota in human metabolic health and disease, Nat Rev Microbiol 19(1) (2021) 55–71.

[4] N. Bosco, M. Noti, The aging gut microbiome and its impact on host immunity, Genes Immun 22(5-6) (2021) 289–303.

[5] M. Witkowski, T.L. Weeks, S.L. Hazen, Gut Microbiota and Cardiovascular Disease, Circ Res 127(4) (2020) 553–570.

[6] M.R. Aljumaah, U. Bhatia, J. Roach, J. Gunstad, M.A. Azcarate Peril, The gut microbiome, mild cognitive impairment, and probiotics: A randomized clinical trial in middle-aged and older adults, Clin Nutr 41(11) (2022) 2565–2576.

[7] P. Luczynski, K.A. McVey Neufeld, C.S. Oriach, G. Clarke, T.G. Dinan, J.F. Cryan, Growing up in a Bubble: Using Germ-Free Animals to Assess the Influence of the Gut Microbiota on Brain and Behavior, Int J Neuropsychopharmacol 19(8) (2016).

[8] C. Lopez-Otin, M.A. Blasco, L. Partridge, M. Serrano, G. Kroemer, Hallmarks of aging: An expanding universe, Cell 186(2) (2023) 243–278.

[9] V.D. Badal, E.D. Vaccariello, E.R. Murray, K.E. Yu, R. Knight, D.V. Jeste, T.T. Nguyen, The Gut Microbiome, Aging, and Longevity: A Systematic Review, Nutrients 12(12) (2020).

[10] C. Tanes, K. Bittinger, Y. Gao, E.S. Friedman, L. Nessel, U.R. Paladhi, L. Chau, E. Panfen, M.A. Fischbach, J. Braun, R.J. Xavier, C.B. Clish, H. Li, F.D. Bushman, J.D. Lewis, G.D. Wu, Role of dietary fiber in the recovery of the human gut microbiome and its metabolome, Cell Host Microbe 29(3) (2021) 394–407 e5.

[11] T. Yatsunenko, F.E. Rey, M.J. Manary, I. Trehan, M.G. Dominguez-Bello, M. Contreras, M. Magris, G. Hidalgo, R.N. Baldassano, A.P. Anokhin, A.C. Heath, B. Warner, J. Reeder, J. Kuczynski, J.G. Caporaso, C.A. Lozupone, C. Lauber, J.C. Clemente, D. Knights, R. Knight, J.I. Gordon, Human gut microbiome viewed across age and geography, Nature 486(7402) (2012) 222-7.

[12] J.D. Hoffman, I. Parikh, S.J. Green, G. Chlipala, R.P. Mohney, M. Keaton, B. Bauer, A.M.S. Hartz, A.L. Lin, Age Drives Distortion of Brain Metabolic, Vascular and Cognitive Functions, and the Gut Microbiome, Front Aging Neurosci 9 (2017) 298.

[13] M.G. Langille, C.J. Meehan, J.E. Koenig, A.S. Dhanani, R.A. Rose, S.E. Howlett, R.G. Beiko, Microbial shifts in the aging mouse gut, Microbiome 2(1) (2014) 50.

[14] K.A. Scott, M. Ida, V.L. Peterson, J.A. Prenderville, G.M. Moloney, T. Izumo, K. Murphy, A. Murphy, R.P. Ross, C. Stanton, T.G. Dinan, J.F. Cryan, Revisiting Metchnikoff: Age-related alterations in microbiota-gut-brain axis in the mouse, Brain Behav Immun 65 (2017) 20–32.

[15] C.S. Wu, S.D.V. Muthyala, C. Klemashevich, A.U. Ufondu, R. Menon, Z. Chen, S. Devaraj, A. Jayaraman, Y. Sun, Age-dependent remodeling of gut microbiome and host serum metabolome in mice, Aging (Albany NY) 13(5) (2021) 6330–6345.

[16] Y. Li, L. Ning, Y. Yin, R. Wang, Z. Zhang, L. Hao, B. Wang, X. Zhao, X. Yang, L. Yin, S. Wu, D. Guo, C. Zhang, Age-related shifts in gut microbiota contribute to cognitive decline in aged rats, Aging (Albany NY) 12(9) (2020) 7801–7817.

[17] J. Siddharth, A. Chakrabarti, A. Pannerec, S. Karaz, D. Morin-Rivron, M. Masoodi, J.N. Feige, S.J. Parkinson, Aging and sarcopenia associate with specific interactions between gut microbes, serum biomarkers and host physiology in rats, Aging (Albany NY) 9(7) (2017) 1698–1720.

[18] X. Zhang, Y. Yang, J. Su, X. Zheng, C. Wang, S. Chen, J. Liu, Y. Lv, S. Fan, A. Zhao, T. Chen, W. Jia, X. Wang, Age-related compositional changes and correlations of gut microbiome, serum metabolome, and immune factor in rats, Geroscience 43(2) (2021) 709–725.

[19] A. Bitto, T.K. Ito, V.V. Pineda, N.J. LeTexier, H.Z. Huang, E. Sutlief, H. Tung, N. Vizzini, B. Chen, K. Smith, D. Meza, M. Yajima, R.P. Beyer, K.F. Kerr, D.J. Davis, C.H. Gillespie, J.M. Snyder, P.M. Treuting, M. Kaeberlein, Transient rapamycin treatment can increase lifespan and healthspan in middle-aged mice, Elife 5 (2016).

[20] K. Kurup, S. Matyi, C.B. Giles, J.D. Wren, K. Jones, A. Ericsson, D. Raftery, L. Wang, D. Promislow, A. Richardson, A. Unnikrishnan, Calorie restriction prevents age-related changes in the intestinal microbiota, Aging (Albany NY) 13(5) (2021) 6298–6329.

[21] B. Sovran, F. Hugenholtz, M. Elderman, A.A. Van Beek, K. Graversen, M. Huijskes, M.V. Boekschoten, H.F.J. Savelkoul, P. De Vos, J. Dekker, J.M. Wells, Age-associated Impairment of the Mucus Barrier Function is Associated with Profound Changes in Microbiota and Immunity, Sci Rep 9(1) (2019) 1437.

[22] J.H. Shin, Y.H. Park, M. Sim, S.A. Kim, H. Joung, D.M. Shin, Serum level of sex steroid hormone is associated with diversity and profiles of human gut microbiome, Res Microbiol 170(4-5) (2019) 192–201.

[23] K. Yoon, N. Kim, Roles of Sex Hormones and Gender in the Gut Microbiota, J Neurogastroenterol Motil 27(3) (2021) 314–325.

[24] L. Yurkovetskiy, M. Burrows, A.A. Khan, L. Graham, P. Volchkov, L. Becker, D. Antonopoulos, Y. Umesaki, A.V. Chervonsky, Gender bias in autoimmunity is influenced by microbiota, Immunity 39(2) (2013) 400–12.

[25] M.C. Decaroli, V. Rochira, Aging and sex hormones in males, Virulence 8(5) (2017) 545–570.

[26] I.H. Fentie, M.M. Greenwood, J.M. Wyss, J.T. Clark, Age-related decreases in gonadal hormones in Long-Evans rats: relationship to rise in arterial pressure, Endocrine 25(1) (2004) 15–22.

[27] H.H. Huang, R.W. Steger, J.F. Bruni, J. Meites, Patterns of sex steroid and gonadotropin secretion in aging female rats, Endocrinology 103(5) (1978) 1855–9.

[28] M. Muller, I. den Tonkelaar, J.H. Thijssen, D.E. Grobbee, Y.T. van der Schouw, Endogenous sex hormones in men aged 40-80 years, Eur J Endocrinol 149(6) (2003) 583–9.

[29] J.F. Randolph, Jr., M. Sowers, I.V. Bondarenko, S.D. Harlow, J.L. Luborsky, R.J. Little, Change in estradiol and follicle-stimulating hormone across the early menopausal transition: effects of ethnicity and age, J Clin Endocrinol Metab 89(4) (2004) 1555–61.

[30] J.K. Goodrich, J.L. Waters, A.C. Poole, J.L. Sutter, O. Koren, R. Blekhman, M. Beaumont, W. Van Treuren, R. Knight, J.T. Bell, T.D. Spector, A.G. Clark, R.E. Ley, Human genetics shape the gut microbiome, Cell 159(4) (2014) 789–99.

[31] A. Kurilshikov, C. Medina-Gomez, R. Bacigalupe, D. Radjabzadeh, J. Wang, A. Demirkan, C.I. Le Roy, J.A. Raygoza Garay, C.T. Finnicum, X. Liu, D.V. Zhernakova, M.J. Bonder, T.H. Hansen, F. Frost, M.C. Ruhlemann, W. Turpin, J.Y. Moon, H.N. Kim, K. Lull, E. Barkan, S.A. Shah, M. Fornage, J. Szopinska-Tokov, Z.D. Wallen, D. Borisevich, L. Agreus, A. Andreasson, C. Bang, L. Bedrani, J.T. Bell, H. Bisgaard, M. Boehnke, D.I. Boomsma, R.D. Burk, A. Claringbould, K. Croitoru, G.E. Davies, C.M. van Duijn, L. Duijts, G. Falony, J. Fu, A. van der Graaf, T. Hansen, G. Homuth, D.A. Hughes, R.G. Ijzerman, M.A. Jackson, V.W.V. Jaddoe, M. Joossens, T. Jorgensen, D. Keszthelyi, R. Knight, M. Laakso, M. Laudes, L.J. Launer, W. Lieb, A.J. Lusis, A.A.M. Masclee, H.A. Moll, Z. Mujagic, Q. Qibin, D. Rothschild, H. Shin, S.J. Sorensen, C.J. Steves, J. Thorsen, N.J. Timpson, R.Y. Tito, S. Vieira-Silva, U. Volker, H. Volzke, U. Vosa, K.H. Wade, S. Walter, K. Watanabe, S. Weiss, F.U. Weiss, O. Weissbrod, H.J. Westra, G. Willemsen, H. Payami, D. Jonkers, A. Arias Vasquez, E.J.C. de Geus, K.A. Meyer, J. Stokholm, E. Segal, E. Org, C. Wijmenga, H.L. Kim, R.C. Kaplan, T.D. Spector, A.G. Uitterlinden, F. Rivadeneira, A. Franke, M.M. Lerch, L. Franke, S. Sanna, M. D’Amato, O. Pedersen, A.D. Paterson, R. Kraaij, J. Raes, A. Zhernakova, Large-scale association analyses identify host factors influencing human gut microbiome composition, Nat Genet 53(2) (2021) 156–165.

[32] E.A. Lopera-Maya, A. Kurilshikov, A. van der Graaf, S. Hu, S. Andreu-Sanchez, L. Chen, A.V. Vila, R. Gacesa, T. Sinha, V. Collij, M.A.Y. Klaassen, L.A. Bolte, M.F.B. Gois, P.B.T. Neerincx, M.A. Swertz, S. LifeLines Cohort, H.J.M. Harmsen, C. Wijmenga, J. Fu, R.K. Weersma, A. Zhernakova, S. Sanna, Effect of host genetics on the gut microbiome in 7,738 participants of the Dutch Microbiome Project, Nat Genet 54(2) (2022) 143–151.

[33] Q. Qi, J. Li, B. Yu, J.Y. Moon, J.C. Chai, J. Merino, J. Hu, M. Ruiz-Canela, C. Rebholz, Z. Wang, M. Usyk, G.C. Chen, B.C. Porneala, W. Wang, N.Q. Nguyen, E.V. Feofanova, M.L. Grove, T.J. Wang, R.E. Gerszten, J. Dupuis, J. Salas-Salvado, W. Bao, D.L. Perkins, M.L. Daviglus, B. Thyagarajan, J. Cai, T. Wang, J.E. Manson, M.A. Martinez-Gonzalez, E. Selvin, K.M. Rexrode, C.B. Clish, F.B. Hu, J.B. Meigs, R. Knight, R.D. Burk, E. Boerwinkle, R.C. Kaplan, Host and gut microbial tryptophan metabolism and type 2 diabetes: an integrative analysis of host genetics, diet, gut microbiome and circulating metabolites in cohort studies, Gut 71(6) (2022) 1095–1105.

[34] M. Hirose, A. Kunstner, P. Schilf, A. Sunderhauf, J. Rupp, O. Johren, M. Schwaninger, C. Sina, J.F. Baines, S.M. Ibrahim, Mitochondrial gene polymorphism is associated with gut microbial communities in mice, Sci Rep 7(1) (2017) 15293.

[35] A. Kunstner, P. Schilf, H. Busch, S.M. Ibrahim, M. Hirose, Changes of Gut Microbiota by Natural mtDNA Variant Differences Augment Susceptibility to Metabolic Disease and Ageing, Int J Mol Sci 23(3) (2022).

[36] T. Yardeni, C.E. Tanes, K. Bittinger, L.M. Mattei, P.M. Schaefer, L.N. Singh, G.D. Wu, D.G. Murdock, D.C. Wallace, Host mitochondria influence gut microbiome diversity: A role for ROS, Sci Signal 12(588) (2019).

[37] R. Sathiaseelan, B. Ahn, M.B. Stout, S. Logan, J. Wanagat, H.V.M. Nguyen, N.G. Hord, A.R. Vandiver, R. Selvarani, R. Ranjit, H. Yarbrough, A. Masingale, B.F. Miller, R.F. Wolf, S.N. Austad, A. Richardson, A Genetically Heterogeneous Rat Model with Divergent Mitochondrial Genomes, J Gerontol A Biol Sci Med Sci 78(5) (2023) 771–779.

[38] E. Bolyen, J.R. Rideout, M.R. Dillon, N.A. Bokulich, C.C. Abnet, G.A. Al-Ghalith, H. Alexander, E.J. Alm, M. Arumugam, F. Asnicar, Y. Bai, J.E. Bisanz, K. Bittinger, A. Brejnrod, C.J. Brislawn, C.T. Brown, B.J. Callahan, A.M. Caraballo-Rodriguez, J. Chase, E.K. Cope, R. Da Silva, C. Diener, P.C. Dorrestein, G.M. Douglas, D.M. Durall, C. Duvallet, C.F. Edwardson, M. Ernst, M. Estaki, J. Fouquier, J.M. Gauglitz, S.M. Gibbons, D.L. Gibson, A. Gonzalez, K. Gorlick, J. Guo, B. Hillmann, S. Holmes, H. Holste, C. Huttenhower, G.A. Huttley, S. Janssen, A.K. Jarmusch, L. Jiang, B.D. Kaehler, K.B. Kang, C.R. Keefe, P. Keim, S.T. Kelley, D. Knights, I. Koester, T. Kosciolek, J. Kreps, M.G.I. Langille, J. Lee, R. Ley, Y.X. Liu, E. Loftfield, C. Lozupone, M. Maher, C. Marotz, B.D. Martin, D. McDonald, L.J. McIver, A.V. Melnik, J.L. Metcalf, S.C. Morgan, J.T. Morton, A.T. Naimey, J.A. Navas-Molina, L.F. Nothias, S.B. Orchanian, T. Pearson, S.L. Peoples, D. Petras, M.L. Preuss, E. Pruesse, L.B. Rasmussen, A. Rivers, M.S. Robeson, 2nd, P. Rosenthal, N. Segata, M. Shaffer, A. Shiffer, R. Sinha, S.J. Song, J.R. Spear, A.D. Swafford, L.R. Thompson, P.J. Torres, P. Trinh, A. Tripathi, P.J. Turnbaugh, S. Ul-Hasan, J.J.J. van der Hooft, F. Vargas, Y. Vazquez-Baeza, E. Vogtmann, M. von Hippel, W. Walters, Y. Wan, M. Wang, J. Warren, K.C. Weber, C.H.D. Williamson, A.D. Willis, Z.Z. Xu, J.R. Zaneveld, Y. Zhang, Q. Zhu, R. Knight, J.G. Caporaso, Reproducible, interactive, scalable and extensible microbiome data science using QIIME 2, Nat Biotechnol 37(8) (2019) 852–857.

[39] M. Martin, Cutadapt Removes Adapter Sequences from High-Throughput Sequencing Reads, embnet 17(1) (2011) 10–12.

[40] B.J. Callahan, P.J. McMurdie, M.J. Rosen, A.W. Han, A.J. Johnson, S.P. Holmes, DADA2: High-resolution sample inference from Illumina amplicon data, Nat Methods 13(7) (2016) 581–3.

[41] K. Katoh, D.M. Standley, MAFFT multiple sequence alignment software version 7: improvements in performance and usability, Mol Biol Evol 30(4) (2013) 772–80.

[42] M.N. Price, P.S. Dehal, A.P. Arkin, FastTree 2--approximately maximum-likelihood trees for large alignments, PLoS One 5(3) (2010) e9490.

[43] N.A. Bokulich, B.D. Kaehler, J.R. Rideout, M. Dillon, E. Bolyen, R. Knight, G.A. Huttley, J. Gregory Caporaso, Optimizing taxonomic classification of marker-gene amplicon sequences with QIIME 2’s q2-feature-classifier plugin, Microbiome 6(1) (2018) 90.

[44] R.R. Rodrigues, M. Gurung, Z. Li, M. Garcia-Jaramillo, R. Greer, C. Gaulke, F. Bauchinger, H. You, J.W. Pederson, S. Vasquez-Perez, K.D. White, B. Frink, B. Philmus, D.B. Jump, G. Trinchieri, D. Berry, T.J. Sharpton, A. Dzutsev, A. Morgun, N. Shulzhenko, Transkingdom interactions between Lactobacilli and hepatic mitochondria attenuate western diet-induced diabetes, Nat Commun 12(1) (2021) 101.

[45] H. Tsugawa, T. Cajka, T. Kind, Y. Ma, B. Higgins, K. Ikeda, M. Kanazawa, J. VanderGheynst, O. Fiehn, M. Arita, MS-DIAL: data-independent MS/MS deconvolution for comprehensive metabolome analysis, Nat Methods 12(6) (2015) 523–6.

[46] A. Agus, J. Planchais, H. Sokol, Gut Microbiota Regulation of Tryptophan Metabolism in Health and Disease, Cell Host Microbe 23(6) (2018) 716–724.

[47] A.B. Larabi, H.L.P. Masson, A.J. Baumler, Bile acids as modulators of gut microbiota composition and function, Gut Microbes 15(1) (2023) 2172671.

[48] H. Lin, Y. An, H. Tang, Y. Wang, Alterations of Bile Acids and Gut Microbiota in Obesity Induced by High Fat Diet in Rat Model, J Agric Food Chem 67(13) (2019) 3624–3632.

[49] Y. Lu, G. Zhou, J. Ewald, Z. Pang, T. Shiri, J. Xia, MicrobiomeAnalyst 2.0: comprehensive statistical, functional and integrative analysis of microbiome data, Nucleic Acids Res 51(W1) (2023) W310–W318.

[50] Z. Pang, G. Zhou, J. Ewald, L. Chang, O. Hacariz, N. Basu, J. Xia, Using MetaboAnalyst 5.0 for LC-HRMS spectra processing, multi-omics integration and covariate adjustment of global metabolomics data, Nat Protoc 17(8) (2022) 1735–1761.

[51] H. Mallick, A. Rahnavard, L.J. McIver, S. Ma, Y. Zhang, L.H. Nguyen, T.L. Tickle, G. Weingart, B. Ren, E.H. Schwager, S. Chatterjee, K.N. Thompson, J.E. Wilkinson, A. Subramanian, Y. Lu, L. Waldron, J.N. Paulson, E.A. Franzosa, H.C. Bravo, C. Huttenhower, Multivariable association discovery in population-scale meta-omics studies, PLoS Comput Biol 17(11) (2021) e1009442.

[52] W. Fusco, M.B. Lorenzo, M. Cintoni, S. Porcari, E. Rinninella, F. Kaitsas, E. Lener, M.C. Mele, A. Gasbarrini, M.C. Collado, G. Cammarota, G. Ianiro, Short-Chain Fatty-Acid-Producing Bacteria: Key Components of the Human Gut Microbiota, Nutrients 15(9) (2023).

[53] S.L. Collins, J.G. Stine, J.E. Bisanz, C.D. Okafor, A.D. Patterson, Bile acids and the gut microbiota: metabolic interactions and impacts on disease, Nat Rev Microbiol 21(4) (2023) 236–247.

[54] S. Nakanishi, T. Serikawa, T. Kuramoto, Slc:Wistar outbred rats show close genetic similarity with F344 inbred rats, Exp Anim 64(1) (2015) 25–9.

[55] A.F. Gileta, C.J. Fitzpatrick, A.S. Chitre, C.L. St Pierre, E.V. Joyce, R.J. Maguire, A.M. McLeod, N.M. Gonzales, A.E. Williams, J.D. Morrow, T.E. Robinson, S.B. Flagel, A.A. Palmer, Genetic characterization of outbred Sprague Dawley rats and utility for genome-wide association studies, PLoS Genet 18(5) (2022) e1010234.

[56] C.S. Carter, A. Richardson, D.M. Huffman, S. Austad, Bring Back the Rat!, J Gerontol A Biol Sci Med Sci 75(3) (2020) 405–415.

[57] S.M. Lee, N. Kim, H. Yoon, R.H. Nam, D.H. Lee, Microbial Changes and Host Response in F344 Rat Colon Depending on Sex and Age Following a High-Fat Diet, Front Microbiol 9 (2018) 2236.

[58] T. Takagi, Y. Naito, R. Inoue, S. Kashiwagi, K. Uchiyama, K. Mizushima, S. Tsuchiya, O. Dohi, N. Yoshida, K. Kamada, T. Ishikawa, O. Handa, H. Konishi, K. Okuda, Y. Tsujimoto, H. Ohnogi, Y. Itoh, Differences in gut microbiota associated with age, sex, and stool consistency in healthy Japanese subjects, J Gastroenterol 54(1) (2019) 53–63.

[59] E. Lkhagva, H.J. Chung, J. Hong, W.H.W. Tang, S.I. Lee, S.T. Hong, S. Lee, The regional diversity of gut microbiome along the GI tract of male C57BL/6 mice, BMC Microbiol 21(1) (2021) 44.

[60] X. Yu, U. Gimsa, L. Wester-Rosenlof, E. Kanitz, W. Otten, M. Kunz, S.M. Ibrahim, Dissecting the effects of mtDNA variations on complex traits using mouse conplastic strains, Genome Res 19(1) (2009) 159–65.

[61] D.A. Merriwether, A.G. Clark, S.W. Ballinger, T.G. Schurr, H. Soodyall, T. Jenkins, S.T. Sherry, D.C. Wallace, The structure of human mitochondrial DNA variation, J Mol Evol 33(6) (1991) 543–55.

[62] M. Cui, A. Trimigno, V. Aru, M.A. Rasmussen, B. Khakimov, S.B. Engelsen, Influence of Age, Sex, and Diet on the Human Fecal Metabolome Investigated by (1)H NMR Spectroscopy, J Proteome Res 20(7) (2021) 3642–3653.

[63] N. Salazar, S. Arboleya, T. Fernandez-Navarro, C.G. de Los Reyes-Gavilan, S. Gonzalez, M. Gueimonde, Age-Associated Changes in Gut Microbiota and Dietary Components Related with the Immune System in Adulthood and Old Age: A Cross-Sectional Study, Nutrients 11(8) (2019).

[64] S. Rahat-Rozenbloom, J. Fernandes, G.B. Gloor, T.M. Wolever, Evidence for greater production of colonic short-chain fatty acids in overweight than lean humans, Int J Obes (Lond) 38(12) (2014) 1525–31.

[65] T.F. Teixeira, L. Grzeskowiak, S.C. Franceschini, J. Bressan, C.L. Ferreira, M.C. Peluzio, Higher level of faecal SCFA in women correlates with metabolic syndrome risk factors, Br J Nutr 109(5) (2013) 914–9.

[66] P. Markowiak-Kopec, K. Slizewska, The Effect of Probiotics on the Production of Short-Chain Fatty Acids by Human Intestinal Microbiome, Nutrients 12(4) (2020).

[67] R. Nagpal, S. Wang, S. Ahmadi, J. Hayes, J. Gagliano, S. Subashchandrabose, D.W. Kitzman, T. Becton, R. Read, H. Yadav, Human-origin probiotic cocktail increases short-chain fatty acid production via modulation of mice and human gut microbiome, Sci Rep 8(1) (2018) 12649.

[68] J.A. Vogt, T.M. Wolever, Fecal acetate is inversely related to acetate absorption from the human rectum and distal colon, J Nutr 133(10) (2003) 3145–8.

[69] L. Drozdowski, T. Woudstra, G. Wild, M.T. Clandindin, A.B. Thomson, The age-associated decline in the intestinal uptake of glucose is not accompanied by changes in the mRNA or protein abundance of SGLT1, Mech Ageing Dev 124(10-12) (2003) 1035–45.

[70] R.P. Ferraris, R.R. Vinnakota, Regulation of intestinal nutrient transport is impaired in aged mice, J Nutr 123(3) (1993) 502–11.

[71] T. Bolin, M. Bare, G. Caplan, S. Daniells, M. Holyday, Malabsorption may contribute to malnutrition in the elderly, Nutrition 26(7-8) (2010) 852–3.

[72] H. Sershen, P. Berger, A.E. Jacobson, K.C. Rice, M.E. Reith, Metaphit prevents locomotor activation induced by various psychostimulants and interferes with the dopaminergic system in mice, Neuropharmacology 27(1) (1988) 23–30.

[73] N. Akram, Z. Faisal, R. Irfan, Y.A. Shah, S.A. Batool, T. Zahid, A. Zulfiqar, A. Fatima, Q. Jahan, H. Tariq, F. Saeed, A. Ahmed, A. Asghar, H. Ateeq, M. Afzaal, M.R. Khan, Exploring the serotonin-probiotics-gut health axis: A review of current evidence and potential mechanisms, Food Sci Nutr 12(2) (2024) 694–706.

[74] Y. Chen, J. Xu, Y. Chen, Regulation of Neurotransmitters by the Gut Microbiota and Effects on Cognition in Neurological Disorders, Nutrients 13(6) (2021).

[75] G. Clarke, S. Grenham, P. Scully, P. Fitzgerald, R.D. Moloney, F. Shanahan, T.G. Dinan, J.F. Cryan, The microbiome-gut-brain axis during early life regulates the hippocampal serotonergic system in a sex-dependent manner, Mol Psychiatry 18(6) (2013) 666–73.

[76] C. Xue, G. Li, Q. Zheng, X. Gu, Q. Shi, Y. Su, Q. Chu, X. Yuan, Z. Bao, J. Lu, L. Li, Tryptophan metabolism in health and disease, Cell Metab 35(8) (2023) 1304–1326.

[77] G. Sui, L. Jia, D. Quan, N. Zhao, G. Yang, Activation of the gut microbiota-kynurenine-liver axis contributes to the development of nonalcoholic hepatic steatosis in nondiabetic adults, Aging (Albany NY) 13(17) (2021) 21309–21324.

[78] S. Comai, C.V. Costa, E. Ragazzi, A. Bertazzo, G. Allegri, The effect of age on the enzyme activities of tryptophan metabolism along the kynurenine pathway in rats, Clin Chim Acta 360(1- 2) (2005) 67–80.

[79] L. Bakker, K. Choe, S. Eussen, I. Ramakers, D.L.A. van den Hove, G. Kenis, B.P.F. Rutten, F.R.J. Verhey, S. Kohler, Relation of the kynurenine pathway with normal age: A systematic review, Mech Ageing Dev 217 (2024) 111890.

[80] J.Y.L. Chiang, J.M. Ferrell, Bile Acid Biology, Pathophysiology, and Therapeutics, Clin Liver Dis (Hoboken) 15(3) (2020) 91–94.

[81] M. Begley, C.G. Gahan, C. Hill, The interaction between bacteria and bile, FEMS Microbiol Rev 29(4) (2005) 625–51.

[82] S. Devkota, E.B. Chang, Interactions between Diet, Bile Acid Metabolism, Gut Microbiota, and Inflammatory Bowel Diseases, Dig Dis 33(3) (2015) 351–6.

[83] S.L. Long, C.G.M. Gahan, S.A. Joyce, Interactions between gut bacteria and bile in health and disease, Mol Aspects Med 56 (2017) 54–65.

[84] H. Suzuki, S. Hayakawa, M. Tajima, Effect of age on plasma bile acids and lipid components in the rat, Lipids 18(9) (1983) 658–60.

[85] G. Lee, H. Lee, J. Hong, S.H. Lee, B.H. Jung, Quantitative profiling of bile acids in rat bile using ultrahigh-performance liquid chromatography-orbitrap mass spectrometry: Alteration of the bile acid composition with aging, J Chromatogr B Analyt Technol Biomed Life Sci 1031 (2016) 37–49.

[86] A. Pena-Cearra, D. Song, J. Castelo, A. Palacios, J.L. Lavin, M. Azkargorta, F. Elortza, M. Fuertes, M.A. Pascual-Itoiz, D. Barriales, I. Martin-Ruiz, A. Fullaondo, A.M. Aransay, H. Rodriguez, N.W. Palm, J. Anguita, L. Abecia, Mitochondrial dysfunction promotes microbial composition that negatively impacts on ulcerative colitis development and progression, NPJ Biofilms Microbiomes 9(1) (2023) 74.

[87] Z. Zhang, H. Zhang, T. Chen, L. Shi, D. Wang, D. Tang, Regulatory role of short-chain fatty acids in inflammatory bowel disease, Cell Commun Signal 20(1) (2022) 64.

[88] I. Guerrero-Encinas, J.N. Gonzalez-Gonzalez, L. Santiago-Lopez, A. Muhlia-Almazan, H.S. Garcia, M.A. Mazorra-Manzano, B. Vallejo-Cordoba, A.F. Gonzalez-Cordova, A. Hernandez-Mendoza, Protective Effect of Lacticaseibacillus casei CRL 431 Postbiotics on Mitochondrial Function and Oxidative Status in Rats with Aflatoxin B(1)-Induced Oxidative Stress, Probiotics Antimicrob Proteins 13(4) (2021) 1033–1043.

[89] M.A. Pascual-Itoiz, A. Pena-Cearra, I. Martin-Ruiz, J.L. Lavin, C. Simo, H. Rodriguez, E. Atondo, J.M. Flores, A. Carreras-Gonzalez, J. Tomas-Cortazar, D. Barriales, A. Palacios, V. Garcia-Canas, A. Pellon, A. Fullaondo, A.M. Aransay, R. Prados-Rosales, R. Martin, J. Anguita, L. Abecia, The mitochondrial negative regulator MCJ modulates the interplay between microbiota and the host during ulcerative colitis, Sci Rep 10(1) (2020) 572.

