## Supplementary Figures for "Age, sex, and mitochondrial-haplotype influence gut microbiome composition and metabolites in a genetically diverse rat model"

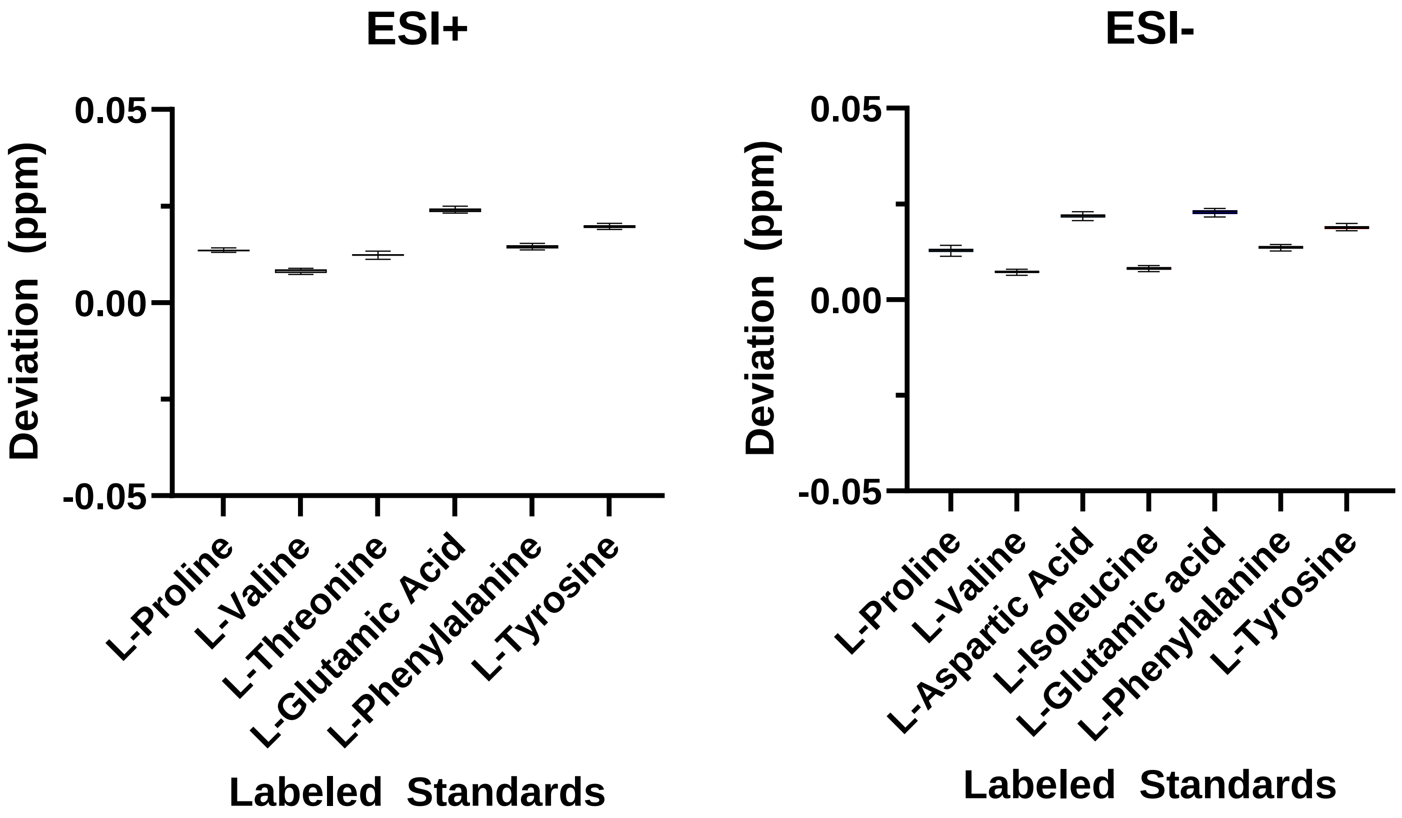


**Supplementary Figure 1: Error deviation for isotopically labeled standards for metabolomic analysis.** Error deviation (ppm) for six isotopically labeled standards (metabolite Mix 1 QReSS Kit (Cambridge Isotope Labs) spiked in our samples (n=61). Deviation error is calculated from the difference between the measured and expected exact mass (m/z value) in both positive (ESI+) and negative (ESI-) electrospray ionization modes. Here we show that our deviation error was <0.05 ppm in all our samples.

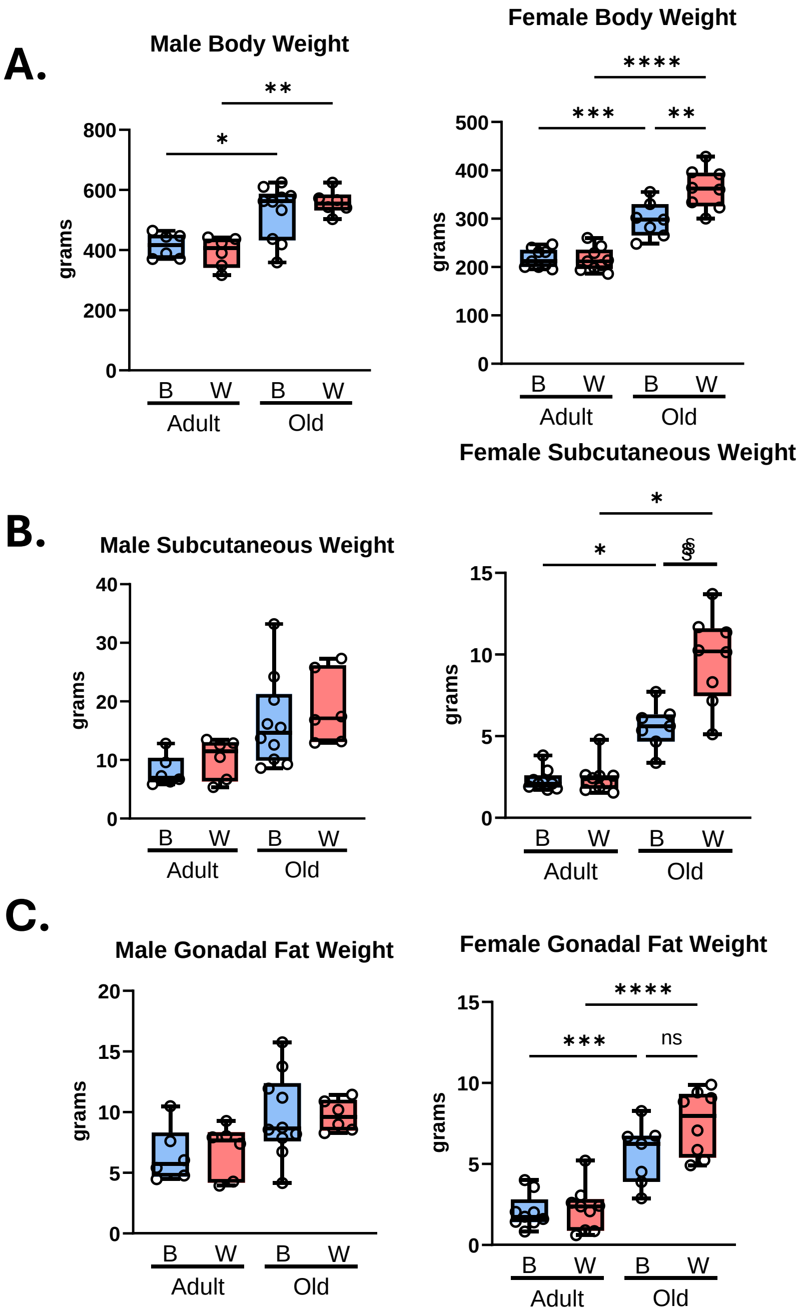


**Supplementary Figure 2. Body composition of male and female OKC-HET$^{B/W}$ rats by age and mt-haplotype.** Body weight (A), subcutaneous fat pad weight (B), and gonadal fat pad weight (C) of male and female, adult (9-months) or old (26-months), and OKC-HET$^{B}$ (blue boxes) and OKC-HET$^{W}$ (red boxes) rats are shown. The box plots display the 1$^{st}$ and 3$^{rd}$ quartiles with the horizontal line for the median, and the whiskers display minimum and maximum values. The data were collected from 6 to 10 rats per group, and statistical significance determined by one-way ANOVA with Tukey's post-hoc ($p<0.05$*, $p<0.01$** $p<0.001$***, $p<0.0001$****) or by the t-test $p<0.05$ $^{\S}$.

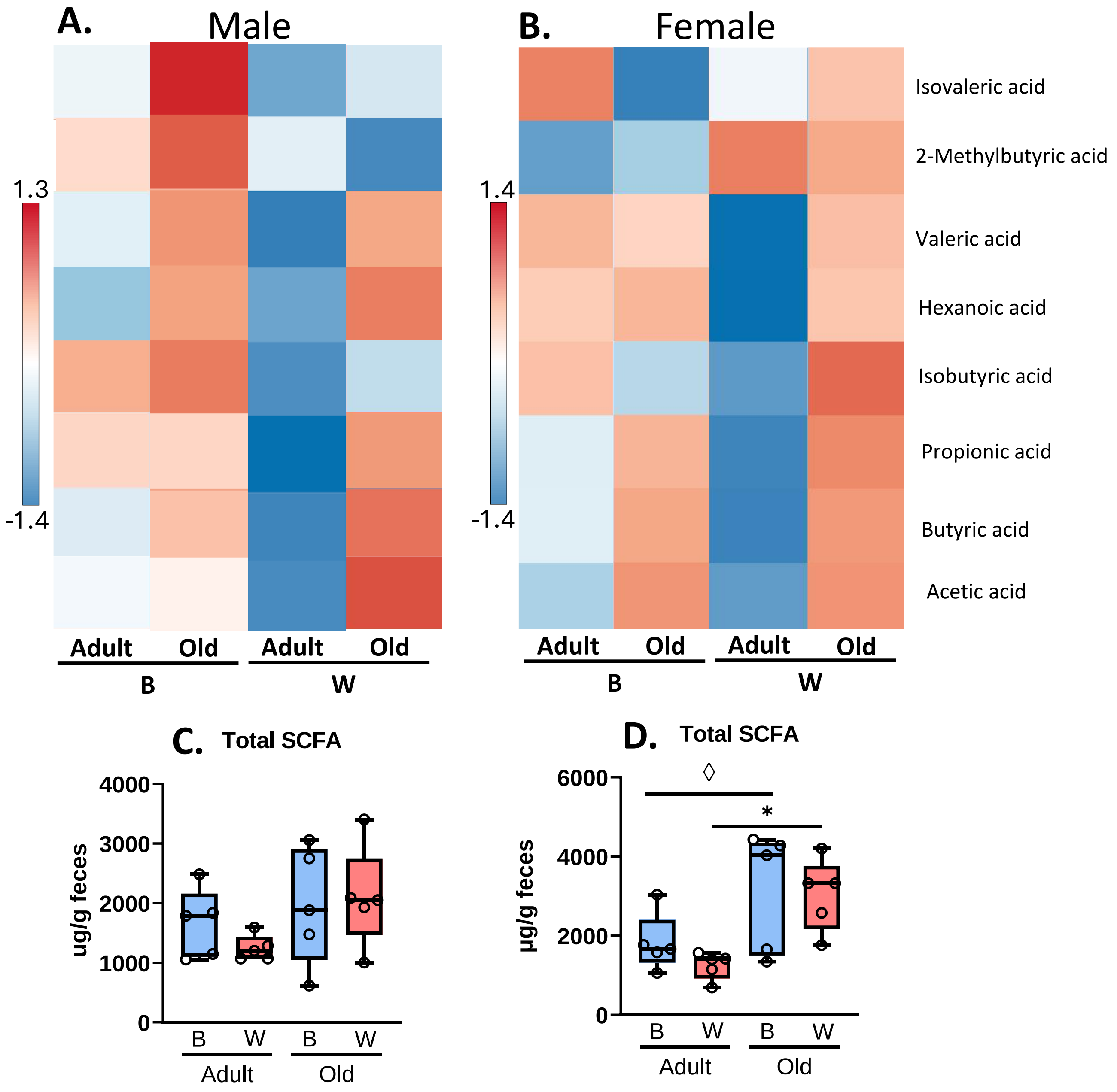


**Supplementary Figure 3. Fecal SCFA profile of OKC-HET$^{W}$ rats.** The heatmap of fecal SCFAs from male (A) and female (B), adult (9-months) and old (26-months) is shown. The total levels of fecal SCFAs are shown for male (C) and female (D) OKC-HET$^{B}$ (blue boxes) and OKC-HET$^{W}$ (red boxes) rats. The boxes display the 1$^{st}$ and 3$^{rd}$ quartiles with the horizontal line for the median, and the whiskers display minimum and maximum values. The data were collected from 5 randomly selected animals per group, and significance was defined as FDR $q<0.05$* or values marginally significance by Fischer's LSD $p < 0.05^{\lozenge}$.

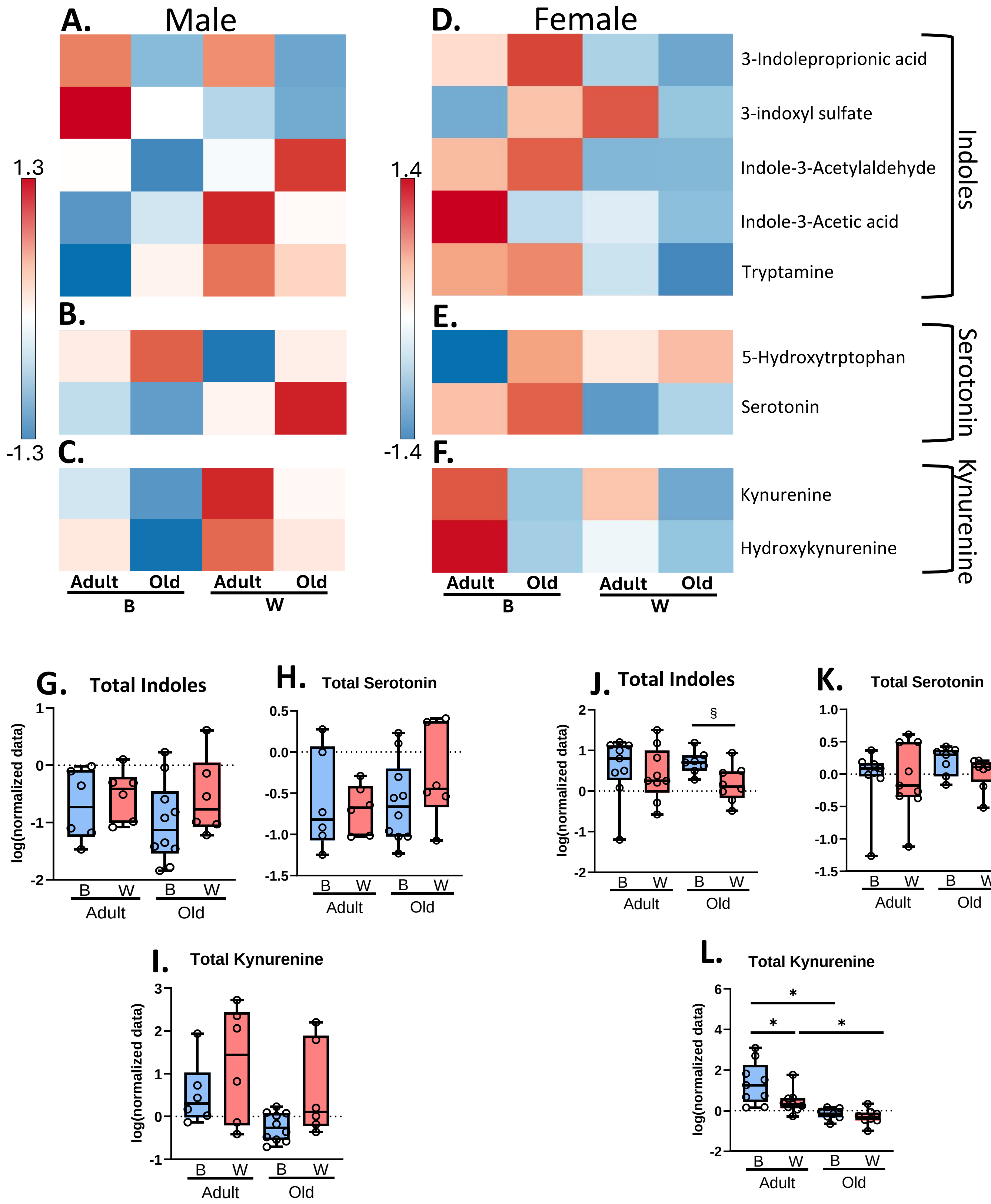


**Supplementary Figure 4. Plasma tryptophan metabolite profile of OKC-HET$^{W}$ rats.** Tryptophan derived metabolites were measured using untargeted metabolomics in the plasma of male and female, adult (9-months) and old (26-months), and OKC-HET$^{B}$ (blue boxes) and OKC-HET$^{W}$ (red boxes) rats. Heatmaps are shown for metabolites of indole (A), serotonin (B), and kynurenine (C) from males and indole (D), serotonin (E), and kynurenine (F) metabolites from females. The total levels of indole metabolites, serotonin metabolites, and kynurenine metabolites are shown for male (G,H,I) and female (J,K,L) rats. The box plots display the 1$^{st}$ and 3$^{rd}$ quartiles with the horizontal line for the median, and the whiskers display minimum and maximum values. The data were collected from 6 to 10 rats per group, and significance was defined as FDR q<0.05* or by the t-test p<0.05$^{§}$.

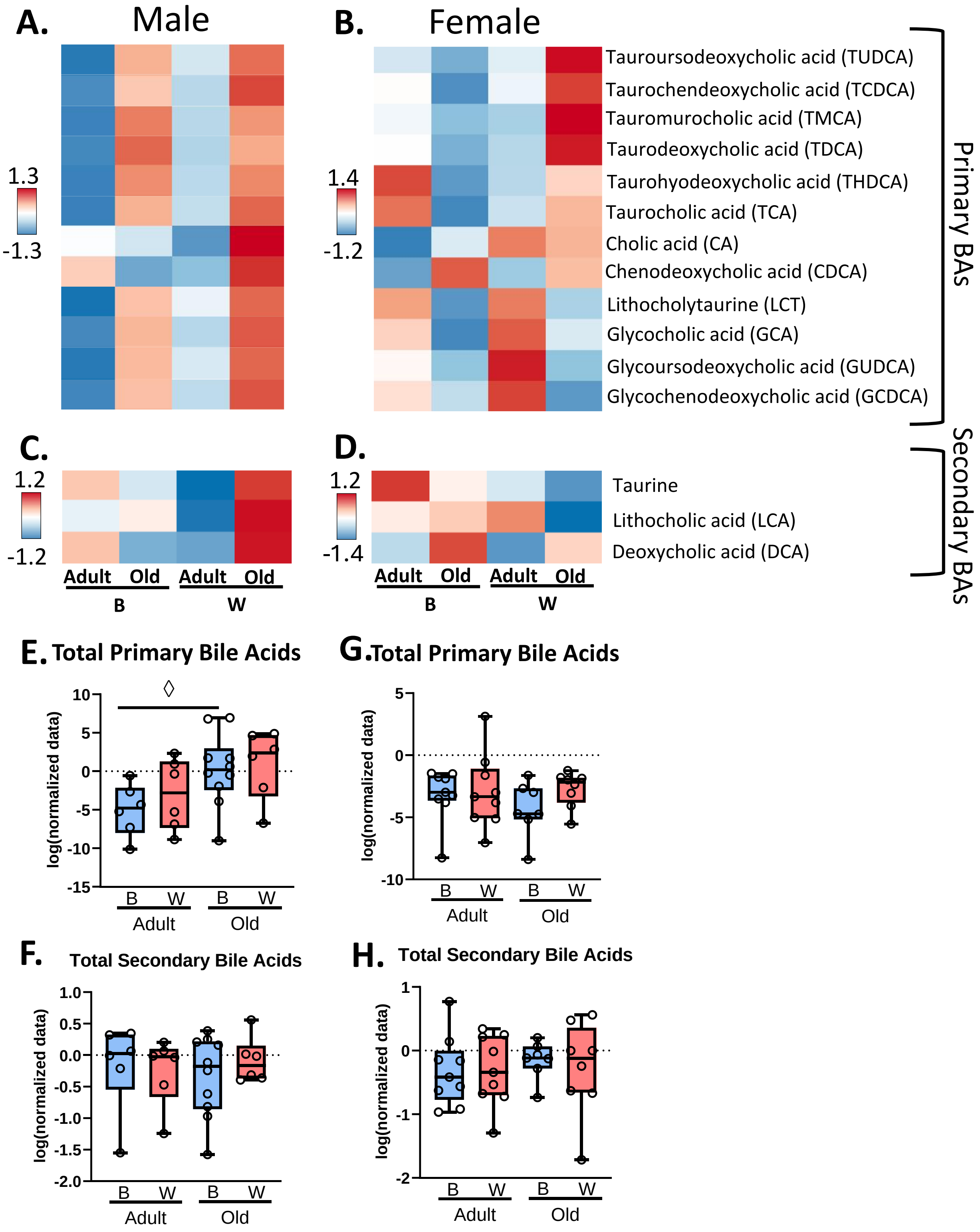


**Supplementary Figure 5. Primary bile acid profile of OKC-HET$^{W}$ rats.** Bile acids were measured using untargeted metabolomics in the plasma of male and female adult (9-months) and old (26-months), and OKC-HET$^{B}$ (blue boxes) and OKC-HET$^{W}$ (red boxes) rats. Heatmaps of primary bile acids for male (A) and female (B) rats and secondary bile acids for male (C) and female (D) rats are shown. The total levels of primary and secondary bile acids from male (E,F) and female (G,H) rats are shown. The box plots display the 1$^{st}$ and 3$^{rd}$ quartiles with the horizontal line for the median, and the whiskers display minimum and maximum values. The data were collected from 6 to 10 rats per group, and $^{\Diamond}$values marginally significant by the Fischer's LSD $p < 0.0^{5}$ shown.

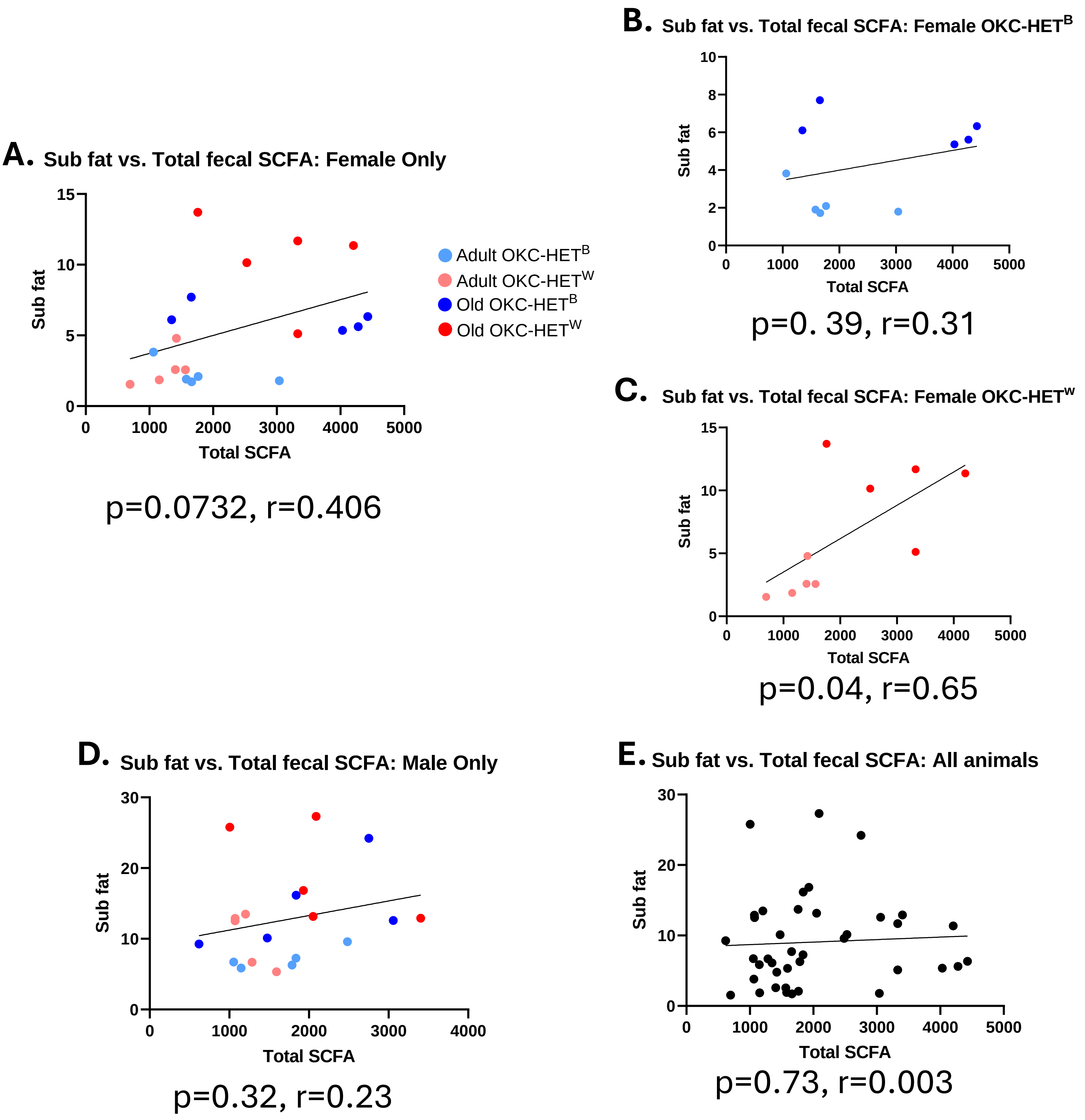


**Supplementary Figure 6. Subcutaneous fat and total fecal SCFA are positively correlated for female OKC-HET$^{W}$ rats.** The correlation between subcutaneous fat (from Supplementary Figure 1) and total fecal SCFAs (from Supplementary Figure 2) are shown for the following: all female rats (A), female OKC-HET$^{B}$ rats (B), female OKC-HET$^{W}$ rats (C), all male rats (D), and all male and female rats (E). The data were collected from 6 to 10 rats per group and statistically analyzed by Pearson's correlation to obtain an r-value. Significance was defined as $p<0.05$.
